# Evidence of a Functionally Segregated Pathway from Lateral Superior Olive to Inferior Colliculus

**DOI:** 10.1101/510354

**Authors:** Nathaniel T. Greene, Kevin A. Davis

**Author notes:** Send Correspondence to: Nathaniel Greene, PhD, Department of Otolaryngology, Research Complex 1-N, Rm 7104, 12800 East 19th Ave, Aurora, CO 80045.

## Abstract

Neurons in the central nucleus of the inferior colliculus (ICC) of decerebrate cats show three major response patterns when tones of different frequencies and levels are presented to the contralateral ear. The frequency response maps of type I units uniquely exhibit a narrowly tuned I-shaped area of excitation around best frequency (the most sensitive frequency) and flanking regions of inhibition at lower and higher frequencies. Type I units receive ipsilateral inhibition, and show binaural excitatory/inhibitory interactions. Lateral superior olive (LSO) principal cells display a similar receptive field organization and sensitivity to interaural level differences (ILDs) and project directly to the ICC, therefore are supposed to be the dominant source of excitatory input for type I units. To test this hypothesis, the responses of ICC units were compared before and after reversible inactivation of the LSO by injection of the non-specific excitatory amino-acid antagonist kynurenic acid. When excitatory activity within the LSO was blocked, many ICC type I units (~50%) were silenced or showed substantially decreased activitycomparable. By contrast, the responses of the other two ICC unit types were largely unaffected. With regard to the origins of unaffected ICC type I units, evidence indicates that the LSO was inactivated in an incomplete, anisotropic manner, and the monaural and binaural responses of such units are similar to those of affected type I units. Taken together, these results support the interpretation that most type I units are the midbrain components of a functionally segregated ILD processing pathway initiated by the LSO.

## Introduction

Single units in the central nucleus of the inferior colliculus (ICC) of unanesthetized decerebrate cats can be grouped into three major types based on the patterns of excitation and inhibition evoked by tones of different frequencies and levels presented to the contralateral ear (Ramachandran et al. 1999). The frequency response maps of type I units uniquely show a narrowly tuned I-shaped area of excitation at frequencies around best frequency (BF; the frequency showing maximum response at approximate 10 dB above threshold) and flanking regions of inhibition at lower and higher frequencies. Type V units produce response maps that exhibit a broad V-shaped excitatory area centered on BF with no signs of inhibition, and type O units show maps that are dominated by inhibition except for an O-shaped island of excitation at frequencies near BF and sound levels near threshold. When tested with dichotic stimuli, with interaural level and time differences (ILDs and ITDs), type I units exhibit binaural excitatory/inhibitory (EI) properties, type V units show binaural facilitation and type O units display only weak effects (Davis et al. 1999; Ramachandran et al. 1999).

The existence of ICC unit types with distinct receptive field properties has led to the proposal that ICC response types are derived from different sources of ascending input that remain functionally segregated within the midbrain (Ramachandran et al. 1999). In particular, it has been speculated that type I units receive their primary excitatory input from principal cells in the lateral superior olive (LSO) (Caird and Klinke 1983; Greene et al. 2010), type V units are shaped by inputs from the medial superior olive (MSO) (Goldberg and Brown 1969; Guinan et al. 1972a; b; Yin and Chan 1990), and that type O units reflect inputs from the dorsal cochlear nucleus (DCN) (Spirou and Young 1991; Young and Brownell 1976). Consistent with this connectionist model, inputs to the ICC form distinct nucleotopic synaptic domains (Oliver and Huerta 1992), unit classification of ICC neurons is not strongly modified by local pharmacological manipulations (Davis 1999; 2002), and each ICC unit type shares the binaural response properties of its putative input (Davis 2002; Ramachandran and May 2002). More recently, Davis (2002) found that most type O units are indeed silenced when the output of the DCN is disrupted. Direct evidence with respect to the other links in this parallel processing model is lacking.

Level tolerant excitatory responses for contralateral tones, flanking inhibitory responses, and binaural EI interactions have been cited as evidence that type I units receive ascending inputs from the LSO (Davis 1999; Ramachandran et al. 1999). Multiple lines of evidence, however, suggest that such properties may be created de novo in the IC or inherited from other non-olivary sources. Consistent with the former interpretation, pharmacological studies reveal that inhibitory inputs shape the frequency tuning and binaural sensitivity of many ICC units (Burger and Pollak 2001; Faingold et al. 1993; LeBeau et al. 2001; Li and Kelly 1992a; b; Vater et al. 1992; Yang et al. 1992). In particular, local blockade of these inputs can result in ICC units showing broader excitatory tuning curves, a loss of sideband inhibition, or loss of binaural suppression (Klug et al. 1995; Park and Pollak 1993). Similarly, intracellular recordings suggest a role for both excitation and inhibition in shaping ILD sensitivity in the bat (Li et al. 2010; Xie et al. 2008; Xie et al. 2007) and cat (Kuwada et al 1997) ICC. Consistent with either de novo creation or a non-LSO source for ILD sensitivity in the ICC, some ICC units retain their EI sensitivity following unilateral and bilateral lesions of the superior olivary complex (Li and Kelly 1992a; Sally and Kelly 1992).

The goal of the present study was to test directly the hypothesis the contralateral LSO projection into the ICC is excitatory, and that this projection provides the dominant excitatory input to ICC type I units. The strength and selectivity of LSO inputs to ICC unit types was assessed by comparing ICC unit responses before and after injection of the reversible excitatory amino acid antagonist kynurenic acid (KYNA) into the LSO. The results reveal that about one-half of ICC type I units are strongly suppressed by LSO inactivation. By contrast, type V and type O units are largely unaffected. With regard to the origins of unaffected ICC type I units, evidence suggests that the LSO was inactivated in an incomplete, anisotropic manner, and analyses of the monaural and binaural responses of affected and unaffected type I units reveal few differences. Taken together, these results support the conceptual model that most ICC type I units are the midbrain components of a functionally segregated ILD processing pathway initiated by the LSO.

## Methods

All surgical and recording procedures in this report were approved by the University Committee on Animal Resources at the University of Rochester.

### Surgical procedures

Details of the surgical preparation are described elsewhere (Greene et al. 2010). Briefly, adult male cats (3-4 kg) with infection-free ears were anesthetized with ketamine (initial/supplemental dose 40/20 mg/kg im) and xylazine (0.5/0.25 mg/kg im), and given atropine (0.05 mg/kg im) to minimize respiratory secretions and dexamethasone (2 mg/kg im) to reduce cerebral edema. Thereafter, core body temperature was maintained at 39°C with a feedback-controlled heating blanket. A catheter was fixed in the cephalic vein to allow infusion of fluids, and a tracheotomy was performed to facilitate quiet breathing. The skin and temporalis muscles overlying the top of the skull were reflected and a craniotomy was made over the left parietal cortex. Cats were made decerebrate by aspirating through the thalamus under visual control; anesthesia was then discontinued. Both ear canals were transected near the tympanic membrane to accept hollow ear bars for delivering closed-field acoustic stimuli. The cat’s head was secured in a stereotaxic apparatus. The left IC was accessed by making a fenestration over the occipital cortex, aspirating the underlying cortical tissue and removing a portion of the bony tentorium. The right LSO was accessed by opening the skull along the midline at the nuccal ridge, and aspirating the cerebellum overlying and bordering the floor of the fourth ventricle caudal to the cerebellar peduncle. At the end of experiments, cats were euthanized with an injection of sodium pentobarbital (100 mg/kg iv).

### Recording protocol

Electrophysiological recordings were made inside a double-walled sound-attenuating chamber (IAC). Acoustic stimuli were delivered bilaterally via electrostatic speakers (TDT) that were coupled to hollow ear bars. These closed-field acoustic systems were calibrated in situ with a probe tube microphone before each experiment, and the resulting functions decreased roughly monotonically from 110 dB SPL at 800 Hz to 90 dB SPL at 48 kHz. Interaural crosstalk was at least 30 dB (and usually > 50 dB) down at all frequencies in the ear opposite to the sound source (Davis 2005; Gibson 1982). All test stimuli, including tones and broadband noise, were created digitally with TDT System 3 hardware. Analog signals were created by passing the waveforms through a 16-bit D/A converter at a sampling rate of 100 kHz. Most stimuli were 200 ms in duration and presented at a rate of 1 burst/s; frequency response maps were constructed from responses to tone bursts that were 50 ms in duration and presented at a rate of 4 bursts/s. All stimuli were gated on and off with 10 ms rise/fall times. Tones were attenuated relative to the acoustic ceiling at each frequency; noise stimuli were flat at the tympanic membrane (i.e., corrected for non-flat calibration curves) and attenuated relative to the maximum spectrum level achievable without any attenuation (~45 dB SL).

The activity of single units in the ICC was recorded with glass-coated platinum-iridium electrodes (1-4 ΜΩ) that were advanced using a motor-controlled multi-electrode positioning system (EPS; Alpha-Omega). The electrode signal was amplified (10,000-30,000x) and bandpass filtered (0.3-6 kHz) using a multi-channel processor (MCP; Alpha-Omega). Template matching software (MSD; Alpha-Omega) was used to discriminate action potentials from background activity. Templates for each unit had error histograms with a single well-defined peak indicating good isolation of one waveform; signal-to-noise ratios were always greater than two-to-one and often greater than ten-to-one. Digitized spike trains were created for on- and off-line analyses by recording spike times relative to stimulus onset.

Recordings in the ICC were made before and after injection of the reversible excitatory amino acid antagonist kynurenic acid into the LSO. Placement of the injection electrode was completed using a two-step process. First, to locate the LSO, platinum-iridium metal electrodes were advanced through the brain stem using a standardized dorsoventral approach (Greene et al. 2010) while 50-ms search stimuli were presented to the excitatory ear (ipsilateral for LSO, contralateral for ICC). When the LSO is approached from this direction, unit BFs tend to increase with depth in its medial and lateral limbs (the BFs are higher in the medial limb), but decrease in its central limb (Tsuchitani and Boudreau 1966). The location of the LSO was verified based upon histological confirmation in one early experiment, but usually upon electrode position (3-4 mm lateral from the midline; 5-6 mm below the floor of the fourth ventricle), progression of unit BFs and similarity of single-unit monaural (tuning curve shape and width, maximum and spontaneous discharge rates) and binaural (i.e. EI) response properties to published data (Finlayson and Caspary 1991; Greene et al. 2010; Guinan et al. 1972a; Guinan et al. 1972b; Tollin and Yin 2005; Tsuchitani 1977). Note that a chopping response in the post-stimulus time histogram is *not* a defining characteristic of LSO units in decerebrate cat (Brownell et al. 1979; Greene and Davis 2012), thus was not used to identify LSO units here.

Second, the metal recording electrode was replaced with a multibarrel glass injection electrode (Kation Scientific LLC) once the LSO was located. One barrel of this pipette contained a carbon fiber for recording unit activity; one or two barrels were filled with KYNA (75 mM in physiological saline, pH 9-10; Sigma); and one, the balance, barrel was filled with 1 M NaCl. The drug and balancing barrels were connected via silver/silver chloride wires to microiontophoresis constant current generators (Harvard Apparatus, BH-2) that were used to generate and monitor retention currents (20 nA, electrode negative) and ejection currents (500 nA, electrode positive). In two early experiments, KYNA was injected into LSO via pressure ejection (~5 μL) through a 30 gauge Hamilton syringe with all but the tip insulated with parylene tubing. In all cases, placement of the injection electrode was verified by audiovisually monitoring the local field potential on an oscilloscope and a monitor speaker for appropriate monaural and binaural responses, at which point the electrode was fixed in place.

### Data collection and analysis

After placing the injection electrode in the LSO (where it remained fixed in place throughout the duration of the experiment), recording electrodes were advanced dorsoventrally through the inferior colliculus, while 50-ms search stimuli (tones or noise) were presented at the BF of the background activity. Entry into the ICC was indicated by a reversal from a decreasing to an increasing progression of BFs (Aitkin et al. 1975; Merzenich and Reid 1974) and by the prevalence of ICC response types that were identified in previous experiments (Davis et al. 1999; Greene et al. 2010; Ramachandran et al. 1999). When an ICC unit was isolated, its BF and threshold were determined, and then the following characterization protocol was initiated. Monaural frequency response maps were measured using isofrequency sweeps (over a 60 dB range in intensities low to high in 2 dB steps) at frequencies alternately above and below BF over a three-octave range (centered on the unit’s BF in 0.1 octave steps); each frequency-intensity combination was presented once. Rate-level functions were obtained for BF tones and broadband noise bursts presented to the excitatory ear by sweeping the level of the stimulus over a 100-dB range (low to high in 1 dB steps). For binaural testing, the same BF tone was presented to both ears, but a 40-dB range of ILDs was created by varying the level of the ipsilateral tone relative to a fixed-level (10 dB re threshold) tone in the contralateral ear.

Responses to acoustic stimuli were recorded again after KYNA was iontophoresed (or in a few cases pressure ejected) into the LSO. The time-course and magnitude of the effects of LSO inactivation was measured by recording ICC unit responses to 100-500 repetitions of 200-ms BF-tone bursts, presented 1 burst/s to the excitatory ear at ~10 dB re threshold. Once a baseline rate was observed (typically 25-50 trials) the retention current was turned off and the ejection current turned on. A minimum of 100 s was allowed for the drug concentration and its effect to stabilize; the agent was applied continuously during data acquisition. Frequency response map, rate-level and ILD functions were repeated. Antagonist application was then discontinued and the cell allowed to recover to baseline activity. Recovery was monitored by recording responses to BF tones. Control experiments in which KYNA-free saline was injected were not conducted. Thus, it is possible that the effects of injections are not due to the effects of KYNA, but are due to the applied current, pH, pressure or some other perturbation. For the purposes of this study, however, the cause of the disruption of ascending LSO influences is not of critical importance as long as the effect is localized.

Responses were analyzed in terms of average discharge rates. Stimulus-evoked rates were computed over the final 80% of the stimulus-on interval in order to exclude onset dominated, and thereby capture steady-state unit responses. Spontaneous rates were computed over the last 50% of the stimulus-off interval. Excitatory (inhibitory) responses were defined as those for which the stimulus-evoked rate was at least one standard deviation above (below) the spontaneous discharge rate. For display purposes only, all response functions were smoothed with a 3-bin triangularly weighted moving-average filter.

## Results

The effects of LSO inactivation were studied on 54 ICC units (40 type I and 14 type O) in 13 cats. The responses of an additional three type V units were recorded, and are included in figures, but will not be discussed. The KYNA injection electrode was localized to the LSO based on histological reconstruction of the electrode path and lesion in one experiment, and by stereotaxic and physiological criteria in the other experiments (Greene et al. 2010). The efficacy of the KYNA in the LSO was verified by recording the responses of LSO units before and after injection, and/or by audiovisual monitoring of the local field potential. The frequency distribution, and monaural and binaural response characteristics of the ICC units were similar to published data (Davis et al. 1999; Ramachandran et al. 1999), though the rate of type I unit recordings was higher because they were of primary interest in this study. Effects in the ICC were observed in 11/13 experiments.

### Electrode placement and effects of KYNA in LSO

The LSO in cat is a relatively large S-shaped 3-dimensional structure, with a highly ordered internal structure (Cant 1984). Placement of the KYNA injection electrode to inactivate the LSO was targeted towards the middle limb. The photomicrograph in Fig. 1A shows the reconstructed path (straight line) of the injection electrode in one of the early pressure injection experiments, shown here since most of the electrode path was visible in more rostral brainstem sections. The most ventral extent of an electrolytic lesion created at the site of the pressure injection, at the end of the experiment, is indicated with an open circle (additional damage to the LSO is visible in more rostral brainstem sections). Note that the tissue damage caused by the syringe tip is confined to the LSO and its immediate surrounding area, suggesting that the injection was localized within this region. Iontophoretic injections were used in most experiments in part to minimize damage resulting from the large injection volume, and to allow precise control over ejection timing. The distribution of best frequencies recorded at LSO injection sites (calculated from single or multi unit activity similarly as in ICC, or estimated from audiovisual assessment of the local field potential) across experiments is shown in Fig. 1B, where the height of each gray shaded bar represents the number of ICC units (of any BF) recorded in each experiment (i.e. at each LSO injection site). The best frequency of recorded units or of the background activity at the injection site was between 6 and 18 kHz in most experiments, consistent with localization within the central limb of the LSO (Tsuchitani and Boudreau 1966).

**Fig. 1.**
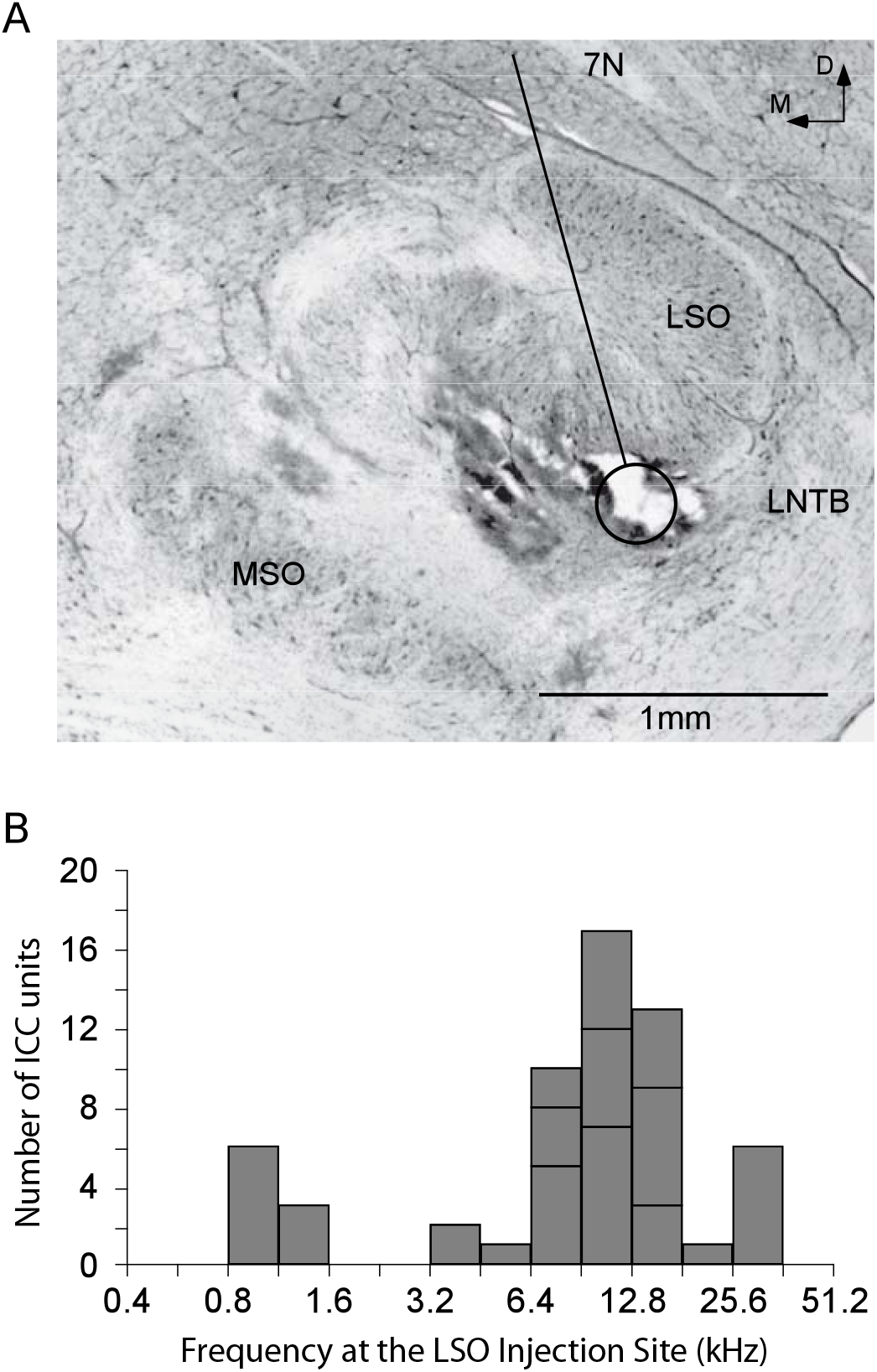
Location and best frequency distribution at the site of kynurenic acid injections. A: The oblique line indicates the path that an insulated 30 gauge Hamilton syringe followed into the LSO (in other slices); the circle indicates the ventral extent of an electrolytic lesion produced at the end of the experiment to aid electrode path reconstruction. The remainder of the electrode tract and the electrolytic lesion are out of this plane of section. Note that the location of the electrolytic lesion suggests that the electrode tip was located within the middle or medial (middle or high BF) limb of the LSO. B: The number of ICC units recorded in each experiment (grouped by gray shaded bars) are shown as a function of the estimated BF at the site of injection in the LSO. M, medial; D, dorsal; MSO, medial superior olive; LNTB, lateral nucleus of the trapezoid body; 7N, seventh cranial (facial) nerve.

The effects of kynurenic acid were studied on several well-isolated, single units within the LSO. The response of a representative LSO unit recorded at the site of injection is shown in Fig. 2A, which shows a spike-time raster (left; each point represents one spike recorded at the time indicated on the x-axis, during the trial indicated on the y-axis), as well as the mean driven discharge rate (right) recorded during each trial. Baseline discharge (for a 200 ms duration tone stimulus) and spontaneous (the final 500 ms of the 1 s recording duration; not shown) rates were recorded for approximately 20-50 seconds (1 trial/sec.) before the injection (gray) to establish baseline firing rates. Injection duration is indicated by a black bar at center. Before injection, the unit was driven strongly by suprathreshold BF-tones presented to the ipsilateral (excitatory) ear (gray). Almost immediately after onset of kynurenic acid injection (black) the response of the unit begins to drop, reaching a new steady-state rate within 1-3 minutes. The onset of the response to each trial is somewhat more resistant to excitatory blockade than the ongoing sustained activity, but was effectively silenced within approximately three minutes. The response of the LSO cell remained suppressed until injection offset, at which point the discharge rate slowly recovered towards baseline (gray).

**Fig. 2.**
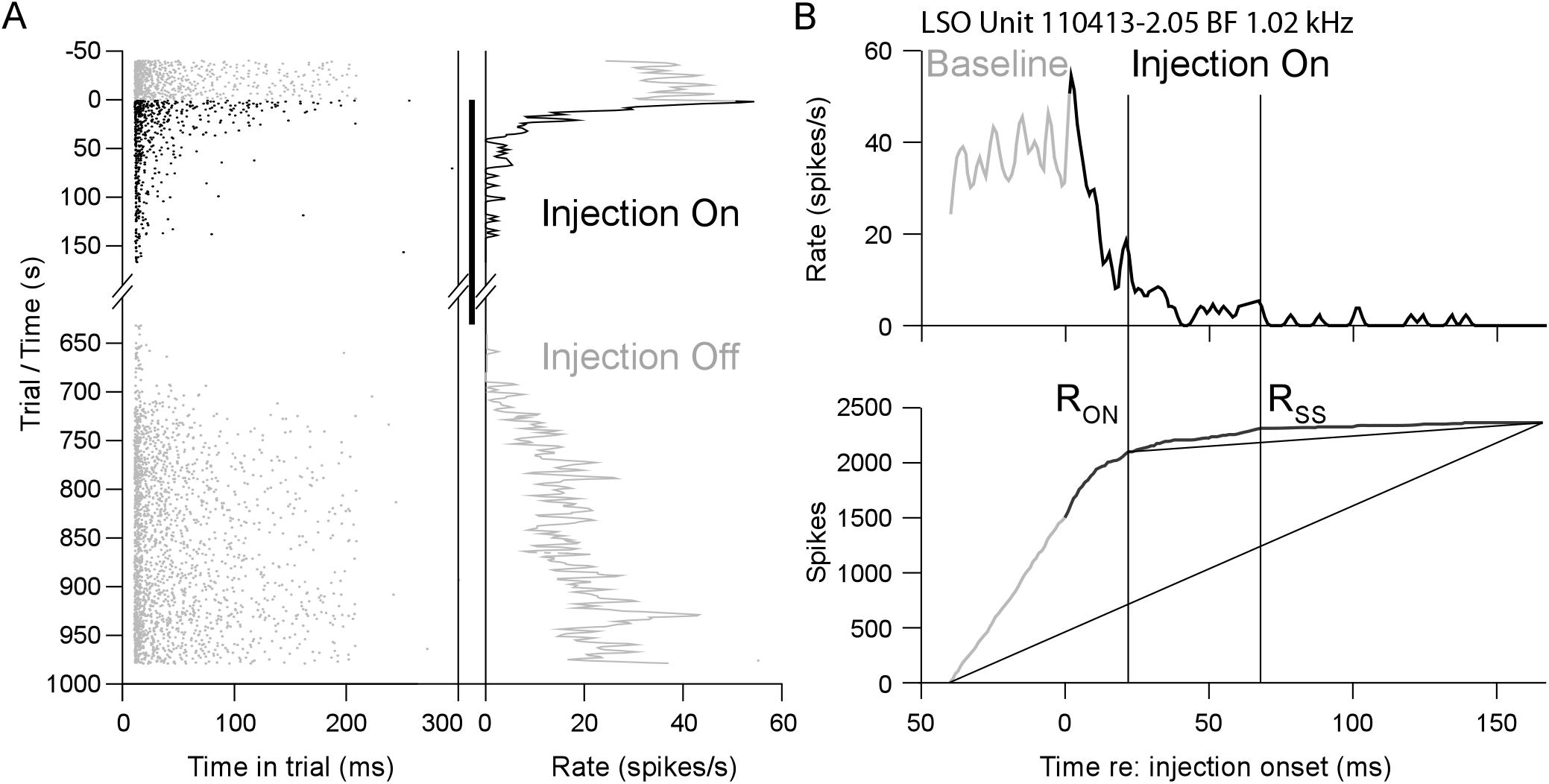
Effects of kynurenic acid at the site of the LSO injection. A: The dot raster (left), and the driven discharge rate (right), which is calculated over the last 80% of the 200 ms stimulus, show the responses of the unit before (gray top) and during (black) kynurenic acid injection, as well as after injection cessation (gray bottom). Each row represents one trial (repeated once per second), and is shown with respect to injection onset (s). The 500 ms gap in recording between the injection trials and the recover trials represents time in which responses to additional stimuli were characterized. B: The latency to the response onset (R_ON_) and steady state (R_SS_) were quantified from the driven discharge rates (top) before (gray) and during (black) injection by calculating the cumulative spike count (bottom). R_ON_ and R_SS_ are defined as the times at which the cumulative deviates maximally from straight lines drawn between the first/R_ON_ and the end of the recording block, respectively.

The latency and magnitude of each unit’s response to KYNA injection were computed using a method modified from Rowland et al. (2007). Mean response to the drug was assessed using the sound-driven discharge rates as a function of time re drug injection (Fig 2B top), from which was calculated the cumulative spike count (Suga et al. 1978) (Fig 2B bottom). Under baseline conditions (gray), LSO (and ICC) units typically respond to repeated tones with a constant rate, thus the cumulative spike count appears as a single straight line. For the example unit shown, KYNA injection resulted in a nearly immediate, profound spike rate decrease, resulting in a decrease in the cumulative spike slope, causing an inflection in the cumulative plot at the onset of the response to the drug. The latencies (vertical lines) to candidate response onsets (R_ON_) following KYNA injection were defined as the points at which the cumulative spike count deviates maximally (vertically) from straight lines drawn between the first and the last points recorded (black diagonal line). The latencies to steady state (R_SS_) were similarly calculated by finding the point on the cumulative spike count that deviated maximally from a line drawn between R_ON_ and the last points recorded (gray diagonal line). The magnitude of the response to KYNA injection was calculated as the difference in mean spike rate between the start of the recording (between the start of recording and R_ON_, i.e. the baseline time period) and the end of the recording (R_SS_ to the last value recorded during the injection; i.e. the injection time period), and is expressed as a percentage change. Statistical significance was evaluated with an unpaired Student’s t-test between the mean discharge rates during the baseline (before R_ON_) and injection (after R_SS_) time periods, and was assessed at p < 0.01. The magnitude of the difference between these two time periods was calculated for all units regardless of whether or not a significant change was noted following KYNA injection. R_ON_ and R_SS_ were not assessed further in units that did not show a significant change. LSO units always showed short-latency, significant decrements in discharge rate following injection of kynurenic acid, as demonstrated by the responses shown in Figs. 2B & 4.

### Effects of LSO inactivation in ICC

In contrast to the fast and powerful responses of LSO units near the site of injection, the responses of ICC units to kynurenic acid injection into LSO were more varied. Figure 3A-C shows the responses (driven and spontaneous discharge rates as a function of time/trial) of three representative type I units before, during and after inactivation of the LSO. Discontinuities in the plots indicate gaps in collection of responses to BF-tones (usually for collection of responses to other stimuli), and KYNA injection was continuous from time zero until the start of the recovery recording (gray). Units can be separated into two groups based on the effect LSO inactivation had on their responses to contralateral (excitatory) BF tones. Unaffected units (Fig. 3A), defined as those units that do not show a significantly different response during the injection than during baseline, show little or no change in response during KYNA injection. Conversely, affected units, i.e. those with a significantly different response during injection, typically show a substantial rate decrease, and can show a somewhat variable R_ON_ latency, i.e. a relatively short (Fig. 3B) or long (Fig. 3C) delay following injection onset. The activity of affected type I units following pressure KYNA injection into LSO decreased in a similar manner as to iontophoretic application (not shown), with responses returning to baseline rates within an hour of injection.

**Fig. 3.**
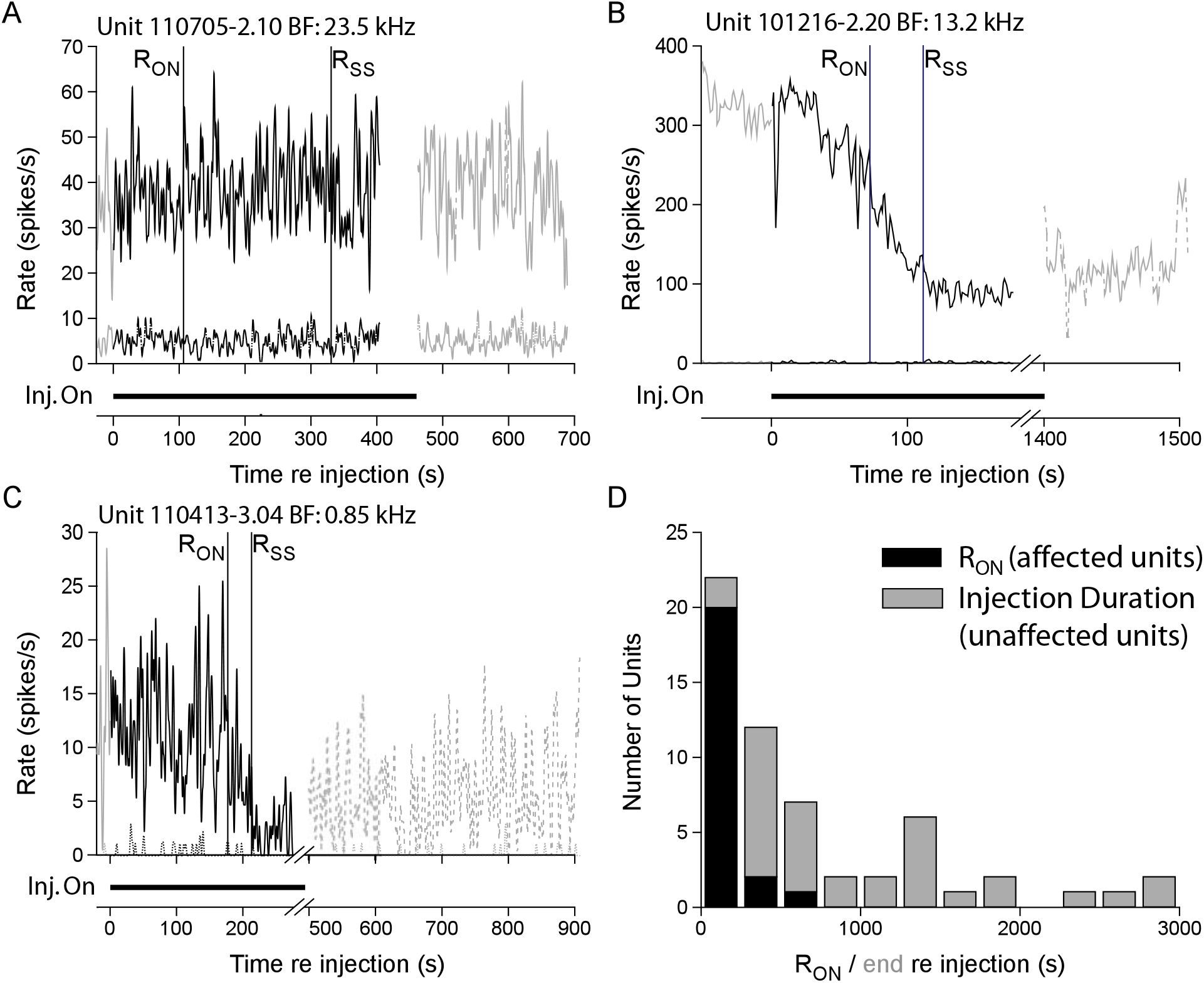
Examples of ICC type I unit responses to LSO inactivation. Type I unit responses could be unaffected (A), or strongly affected with onset delays (R_ON_) varying between relatively short (B; < 100s) or long (C; > 200s). Responses shown are driven and spontaneous discharge rates as a function of time after injection. Effect magnitude was quantified as a percentage change between the time periods before R_ON_ (including before injection onset, black) and after R_SS_ (recovery period excluded). Significance was assessed with a student’s t-test (criterion: p < 0.01). D: Magnitude of the response change (%) is shown as a function of R_ON_ latency for units significantly affected (black), and as a function of total injection duration for unaffected units (gray). Discharge rates were usually assessed for at least 300s, during which most significantly affected units showed a response to the injection. Injection durations were typically much longer if no significant response was observed.

As demonstrated in Fig. 3B-C, R_ON_ latency can vary considerably within the population of significantly affected units. This variability is presumably a result of varying distances (and thereby diffusion times) of LSO projection cells from the site of injection. A histogram of R_ON_ latencies for significantly affected ICC units is shown in Fig. 3D (black bars). For comparison, gray bars show the injection duration for unaffected ICC units. R_ON_ latencies ranged between 25s and 750s with a median R_ON_ latency of just over 100 seconds. Note, all but two affected cells showed a response by 300 seconds after injection onset, whereas injection durations were at least 300 seconds (and often much longer) in all but two unaffected units, suggesting that injection duration was not likely a limiting factor in the observation of unaffected units following LSO inactivation.

The effects of LSO inactivation were different among ICC unit types (i.e. V, I or O). The specificity of LSO projections onto ICC type I units is demonstrated most directly in Fig. 4, which shows data from a type V, I, and O units recorded sequentially in one electrode penetration in one experiment. These units were located at similar recording depths in the ICC and thus had similar BFs. Each ICC unit is shown in conjunction with the response of a single LSO unit recorded simultaneously with all three ICC units (same as in Fig. 2). This LSO unit had a similar BF as the ICC units, and provided a clear indication of the drug’s effect at the site of injection. Injection of KYNA into the LSO dramatically decreased discharge rates in the type I unit (Fig. 4B). By contrast, neither the type V (Fig. 4A) nor the type O (Fig. 4C) units showed a substantial rate change following drug injection despite a comparable suppression of the LSO unit in each case. Note that tone frequencies and intensities were set at the BF, and 20 dB above threshold, of the ICC unit in the *bottom* of each plot, resulting in the slight variability of the LSO unit’s response. The LSO unit’s activity was allowed to recover to baseline (at least 30 minutes) between each set of recordings.

**Fig. 4.**
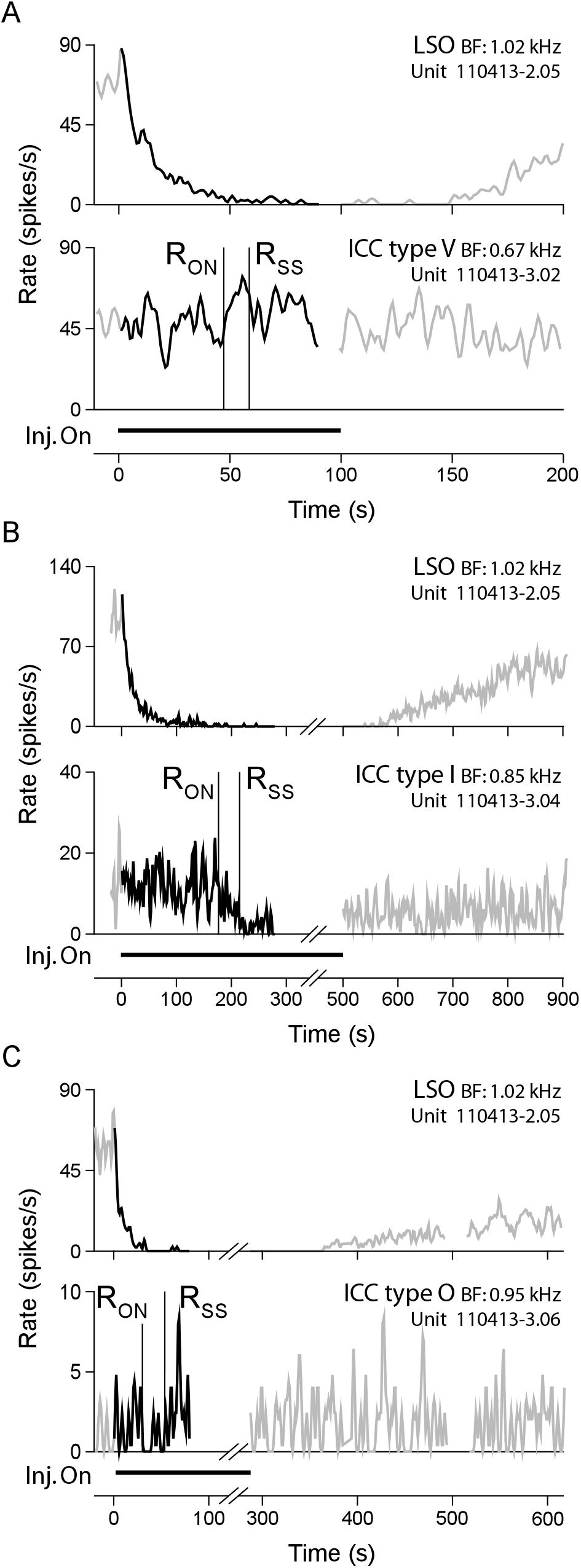
Typical responses of each ICC unit type to kynurenic acid injection into LSO. *Top*, responses of a single LSO unit (same unit as in Fig. 2) were recorded simultaneously with ICC units before (gray left), during (black) and after (gray right) kynurenic acid injection. The type V (A), type I (B), and type O (C) units shown were recorded at a similar depth, and with similar best frequencies, in a single electrode penetration. Sounds were set at the BF and 20 dB above threshold of the ICC unit in the *bottom* of each plot.

Figure 5 shows the magnitude of the effect of LSO inactivation on the different ICC unit types; significantly affected units are shown with shaded bars and unaffected units with hashed white bars. Overall, significantly affected units (23/57) usually showed a 60% or greater rate decrease following KYNA injection (17/23), but could show a rate change magnitude (decrease or increase) of as low as 25%. Unaffected units typically show rate changes less than 20%. With regard to the different ICC unit types, approximately one-half of type I units were strongly suppressed by LSO inactivation (19/40), whereas most type O (13/14) and V (2/3) units were not significantly affected. Two type I units showed small but significant increases in rate (< 50%). The remaining type I (19/40) units show little or no rate change following injection.

**Fig. 5.**
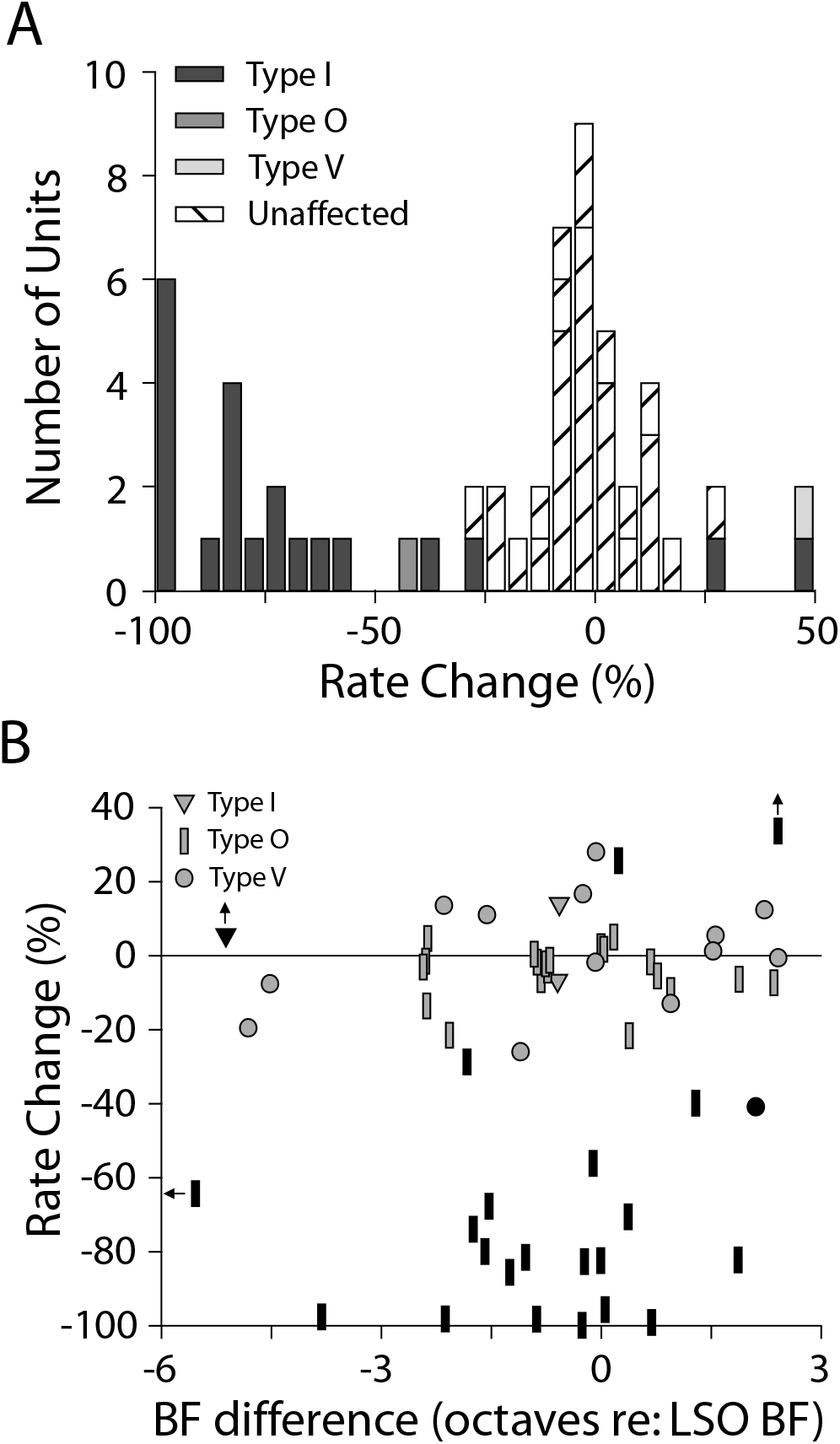
Rate change of all ICC units. A: Histogram of the change in rate (%) is shown for units showing a significant effect of kynurenic acid injection (gray), and for unaffected units (of all types; hashed white bars). B: Rate change (%) for each unit as a function of the difference in that unit’s BF and the estimated BF at the LSO injection site in octaves. Symbol indicates the ICC unit type, and units significantly affected by KYNA injection are indicated with black symbols.

The relationship between the BF at the LSO injection site from recorded single-unit activity or estimated from the local field potential) and the BF of the recorded ICC unit was assessed by plotting the rate change (effect magnitude) as a function of the difference in BFs between the two sites (in octaves re: the BF at the LSO injection site) in Fig. 5B. Most ICC units were recorded within ~ 3 octaves of the LSO injection site BF. Both significantly affected and unaffected units were found across the range of BF differences tested, although significantly affected units were most often recorded within ~1.5 octaves, suggesting a weak BF dependence, and thus a broad (though perhaps not complete) spread of the KYNA injection within the LSO.

### Affected type I units

Basic monaural and binaural response characteristics of ICC units were studied before and after KYNA injection into LSO. Predictably, response characteristics related to discharge rate were altered in affected units, and no differences were observed for unaffected units (not shown). The frequency response map of a representative affected type I unit is shown before (gray), during (black), and after (gray/white dashed) LSO inactivation in Fig. 6A. Each curve shows the unit’s driven discharge rate (height) as a function of frequency at a single sound level (in dB SPL). Spontaneous rates are indicated by horizontal edges on the gray shaded segments, and were consistent across stimulus levels; driven rates above (dark gray) and below (light gray) the spontaneous rate indicate excitation and inhibition respectively. Under control conditions, type I responses show a narrow V shaped excitatory response area at BF (vertical line), with areas of inhibition at higher and lower frequencies. During KYNA injection into LSO, spontaneous (as indicated by the rate suppression at all frequencies) and driven activity is reduced, from a maximum of approximately 55 to 15 spikes/s. While incomplete, the recovery function (dashed) shows a partial return to the baseline discharge rate.

**Fig. 6.**
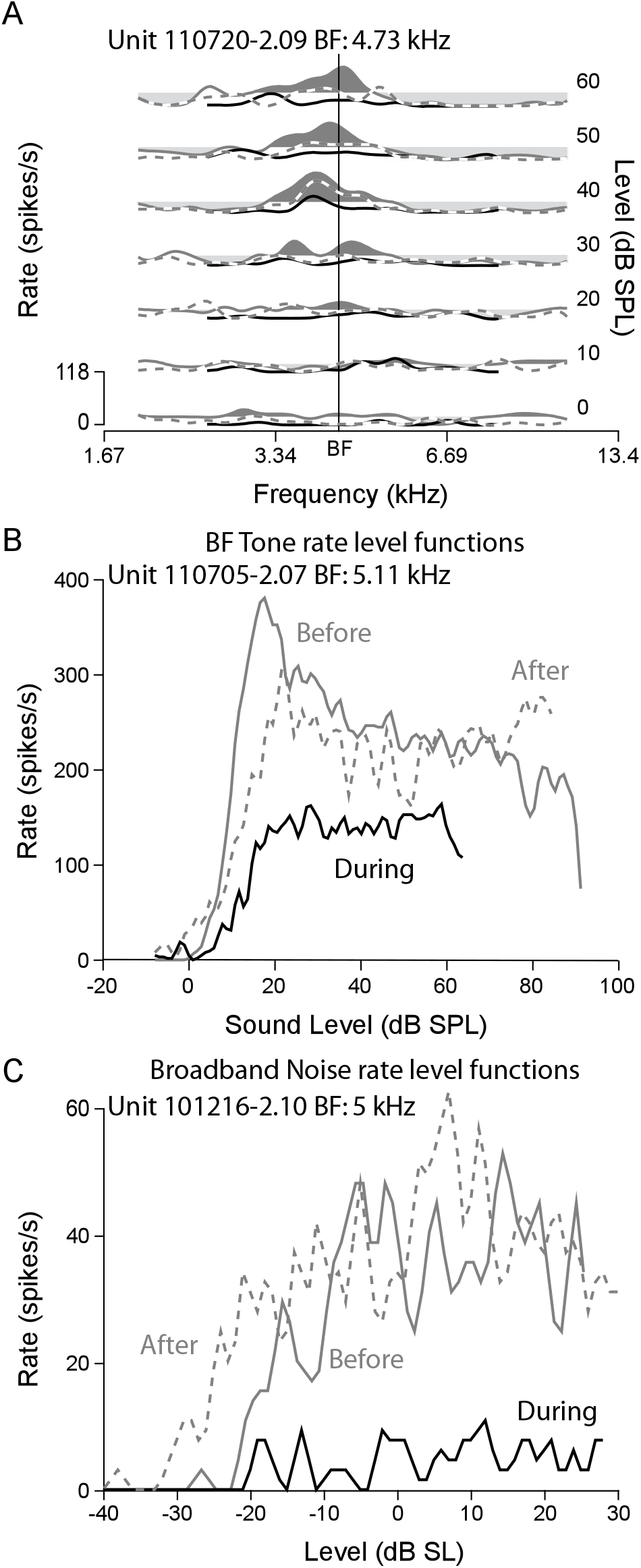
LSO inactivation suppresses the discharge rates of affected type I units, but affects little the frequency and level tuning. Responses shown are from three independent units. A: The frequency response map of an example ICC type I unit (110720-2.09) before (gray shading: dark: responses > spontaneous, light: responses < spontaneous), during (black line) and after (dashed gray/white line) injection of kynurenic acid into LSO. Response rate (height, spikes/s) is shown at several levels (rows, dB SPL) as a function of stimulus frequency (kHz). Best frequency (BF) is represented with a vertical line. BF-tone (B) and broadband noise (C) rate-level functions of affected type I units (B: 110705-2.07; C: 101216-2.10) before (gray) during (black) and after (dashed) kynurenic acid injection. Dark (light) shaded areas represent responses above (below) the baseline spontaneous rate (assessed during the 800ms of quiet following each tone presentation during the baseline recording), representing an excited (inhibited) response.

Example rate-level functions for (different) affected type I units in response to BF-tones and noise are shown in Fig. 6B and 6C, respectively. In these plots, the driven and spontaneous (near zero) discharge rates are shown for control (gray), injection (black), and recovery (gray dashed) conditions. Under control conditions, the responses of type I units climb to a maximum and then either maintain a steady discharge rate at higher stimulus levels (as in the case of the response to noise here; Fig. 6C) or decline to some extent (tones; Fig. 6B). During LSO inactivation the elicited discharge rates decrease, but the basic response characteristics (i.e. threshold, and monotonicity) remain largely unchanged for both stimulus types (Table 1). In two type I units the effects of LSO inactivation were disparate, where the discharge rate increased in response to noise stimuli despite a reduced rate to tones (not shown).

**Table 1:** Affected type I unit response characteristics before and during LSO inactivation. Values are medians across the population of significantly affected type I units. Parameters that show a significant difference (Mann-whitney U test, p < 0.05) are marked with a *.

|  | ICC type I units |  |  |  |
| --- | --- | --- | --- | --- |
|  | Before<br>injection | N | During<br>injection | N |
| BF-tone threshold re ANF (dB) | 6 | 20 | 12.5 | 16 |
| Noise threshold (dB) | -28.5 | 14 | -27 | 13 |
| Q<sub>10</sub> re narrowest ANF | -2.49 | 17 | -2.67 | 8 |
| Q<sub>40</sub> re narrowest ANF | 0.47 | 19 | 0.06 | 11 |
| Spontaneous rate (spikes/s) | 4.59 | 20 | 0.44 | 16 |
| Max rate for BF tones (spikes/s) | * 109.0 | 20 | 13.5 | 16 |
| Max rate for noise (spikes/s) | 27.5 | 13 | 10.7 | 11 |
| Norm. slope for tones (%/dB) | -0.0043 | 20 | -0.0071 | 16 |
| Norm. slope for noise (%/dB) | -0.0089 | 13 | 0 | 11 |
| ILD % inhibition | 51.2 | 12 | 68.7 | 10 |
| ILD max slope | * -1.9 | 12 | -1.22 | 10 |
| ILD50 | 3.12 | 12 | -0.9 | 10 |

Figure 7A demonstrates the typical change that a type I unit ILD function exhibits from before (gray) and during (black) LSO inactivation. ILD curves were generated by fixing the sound level in the contralateral (excitatory) ear at approximately 10 dB above threshold, and increasing the level in the ipsilateral (inhibitory) ear from 20 dB below to 20 dB above this level (+ 20 dB ILD to − 20 dB ILD contra re ipsi), which spans most of the physiologically relevant range of ILD for the cat (Tollin and Koka 2009a; b). Sigmoid functions (dashed) were fit (least squares) to each curve in order to quantify several characteristics (e.g. half-max ILD, percent reduction, maximum slope; mean values across the population of affected units provided in table 1) of the response (Tollin et al. 2008). Before injection this unit produced a characteristic EI ILD sensitivity, that, at increasing ipsilateral (inhibitory) sound levels, elicited a strong suppression of the response towards the spontaneous rate. During injection, the response of this unit showed a similar trend, but the overall rate of the response decreased, and the dynamic range (i.e. the range of driven discharge rates observed under each condition) was reduced. This unit showed no spontaneous activity either before or during KYNA injection. Figure 7B shows the dynamic range during the injection as a function of the dynamic range before the injection for all type I units significantly reduced by LSO inactivation. All units lie below or near the unity line, thus show greater than or equal dynamic range in the baseline condition than injection. Dynamic range, however, can decrease by two mechanisms: a decrease in the excitatory drive resulting in a *reduced* rate at +20 dB ILD, or a decrease in the inhibitory input resulting in an *increased* rate at −20 dB ILD. To differentiate between these two possibilities we found the difference between the baseline and injection sigmoid fits at these two ILD values. The rate decrease (in spikes per second) at −20 dB versus +20 dB ILD is plotted in Fig. 7C. Consistent with the loss of an excitatory, ILD-sensitive input, most units show a greater rate decrease at +20 then −20 dB ILD.

**Fig. 7.**
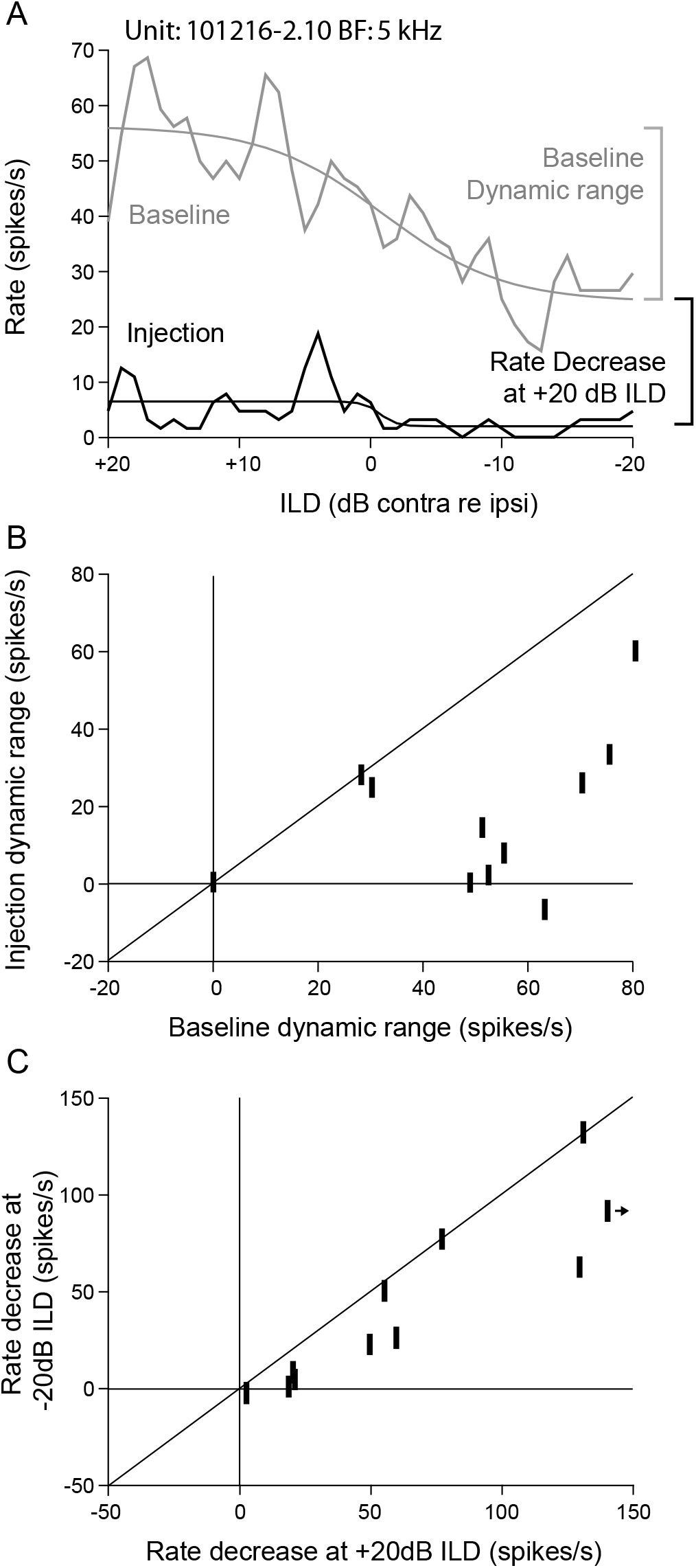
The effects of LSO inactivation on affected type I units is consistent with loss of an ILD sensitive excitatory input. A: ILD functions, i.e. driven discharge rate as a function of difference in level between the two ears at approximately 10 dB re threshold (in dB contra re ipsi sound level), are shown for an example ICC type I unit before (gray) and during (black) injection of kynurenic acid into LSO. Sigmoid fits are shown as smooth lines superimposed on each curve. Bars at right illustrate the definitions of dynamic range (i.e. the driven discharge rate change between +20 and −20 dB ILD *within* the baseline and injection curves), and rate decrease (i.e. the decrease in rate at + 20 dB and at - 20 dB *across* Baseline to Injection). B: The dynamic range of ILD sigmoid fits during the injection are shown as a function of the dynamic range before injection. C: The rate decrease between injection and baseline sigmoid fits at −20 dB ILD (inhibition dominates) are compared to the rate changes at +20 dB ILD (excitation dominates). B and C show only the responses of significantly reduced type I units (defined as in Fig. 6).

LSO inactivation produced more complex results in a minority (3/40) of type I units. The frequency response map shown in Fig. 8A demonstrates the changes in rate produced by LSO inactivation as a function of frequency and level. In the baseline condition (black line/dark gray shading) the type I unit shows a narrow V-shaped excitatory response area (dark gray shaded area) with side-band inhibition (i.e. the solid black line drops below the horizontal black line that represents the spontaneous rate), comparable to the prototypical type I unit (e.g. Fig. 5A). Following LSO inactivation (light gray line), however, the discharge rate is reduced relatively strongly at high sound levels, while responses near threshold show a more moderate reduction. This trend is more clearly visible in the BF-tone rate level functions shown in Fig. 8B. The shape of the BF-tone rate-level curve during KYNA injection (gray) becomes more non-monotonic, and drops below the baseline spontaneous rate (black horizontal line), and nearly to the injection spontaneous rate (gray horizontal line) (. LSO inactivation, therefore, results in a change in response type for this minority of units, from a clear type I to a response approaching that of a type O. The observed effect magnitude as a function of time re: injection is shown in Fig. 8C. Since the time-course of effects following KYNA injection were assessed using BF tones presented at relatively low sound levels, the observed response decrease during injection (~25%; Fig. 8C) somewhat undervalues the discharge rate decrease at higher levels (> 50%) following LSO inactivation.

**Fig. 8.**
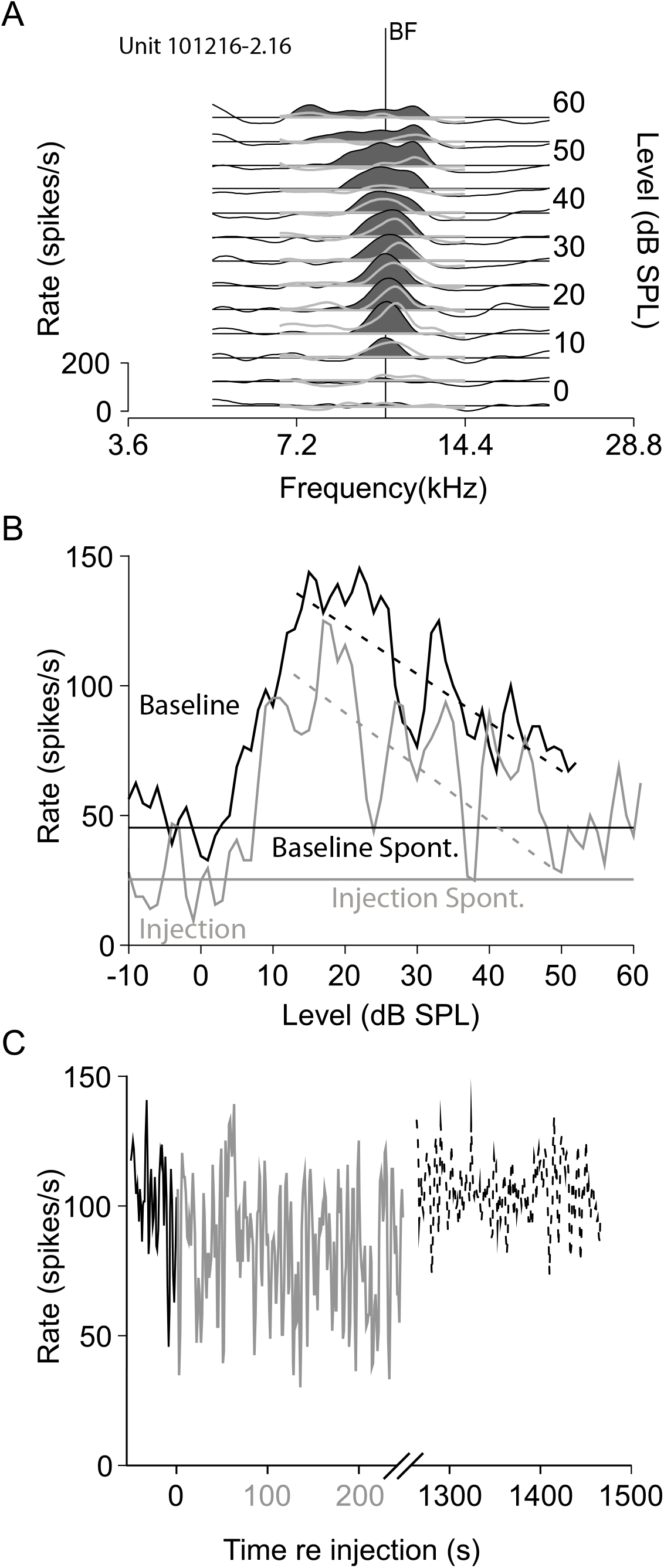
The frequency response map of an example type I unit reveals a change from an I type before to an O type response map during injection of kynurenic acid into LSO (A). BF (10.2 kHz) is indicated with a vertical line. B: BF tone rate-level functions were recorded for baseline (black) and during KYNA injection (gray) conditions. Driven discharge and spontaneous rates are shown. Straight lines (dashed) were fit to the plateau region of the response. C: The response of this unit as a function of time after injection (s), at approximately 20 dB above threshold, reveals a small (~25 %) but *not* significant change during KYNA injection.

### On unaffected type I units

Although LSO inactivation has a strong effect on the discharge rates of many type I units, a large proportion remain unaffected. One potential explanation for this disparity is that KYNA injection simply failed to inactivate completely the LSO. Circumstantial evidence for this possibility is shown in Figure 9, which shows the response maps of sequentially recorded type I units from single electrode penetrations in two experiments, displayed according to BF of the unit relative to the injection site in LSO. In 9A, the units with BFs above and below the LSO BF (~ 5 kHz) showed responses during KYNA injection (dark gray) that were comparable to baseline (light gray), whereas the unit with a similar BF showed strong rate suppression. Conversely, 9B shows an electrode penetration where units with differing BFs showed significantly affected responses, while the unit with a similar BF as LSO (~ 6 kHz) was unaffected. These two seemingly contradictory results might be explainable by differential spread of drug within the LSO: preferentially along an isofrequency contour (and sparing distant isofrequency contours) in the first case, and along the tonotopic axis (sparing distant neurons within the same isofrequency contour) in the second. Further explanation of these two possibilities is provided in the discussion.

**Fig. 9.**
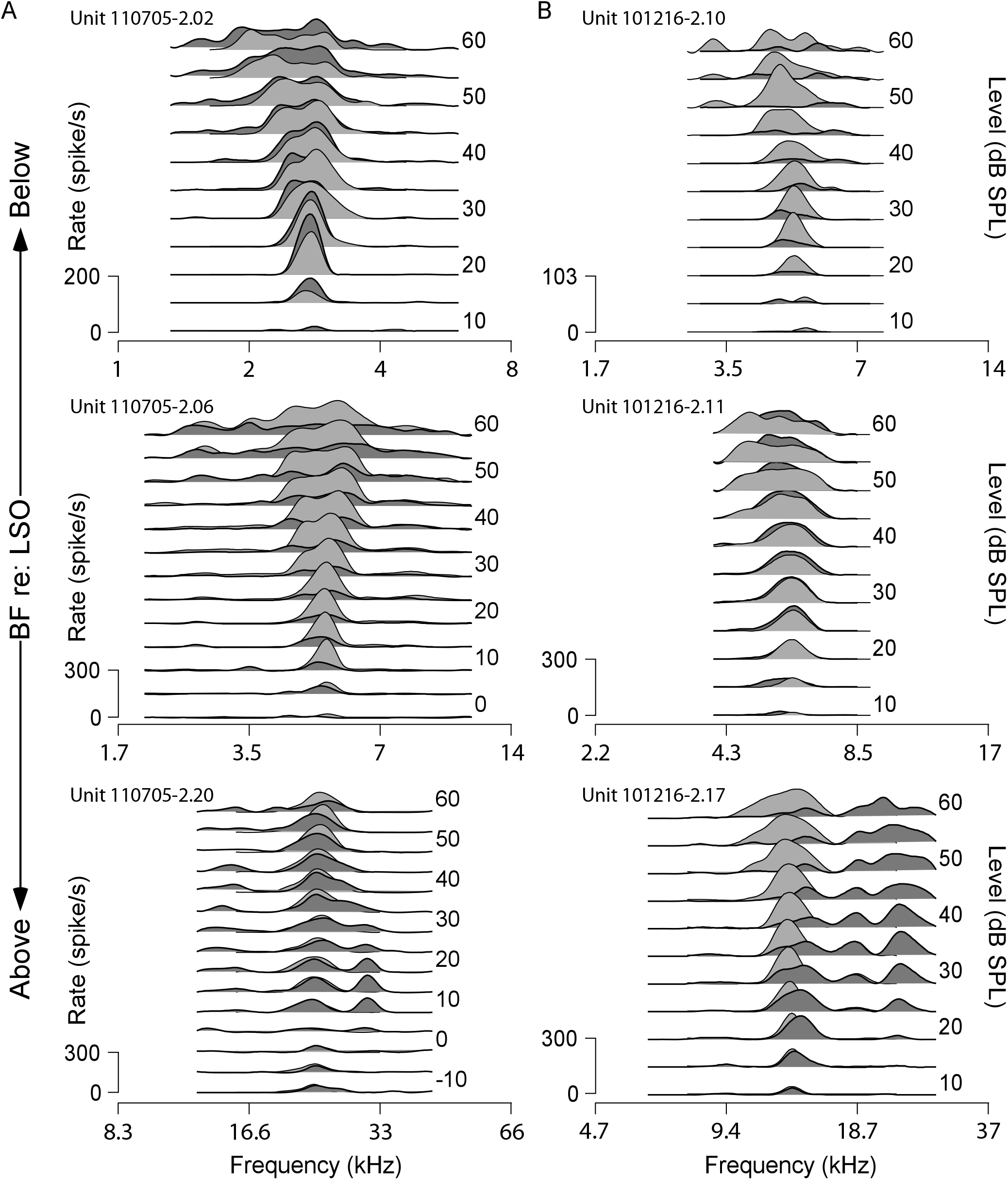
Frequency response maps of ICC type I units that were sequentially recorded, in single electrode penetrations, in two experiments A: 110705, and B: 101216, reveal affected units with similar (A) and dissimilar (B) frequencies as the LSO injection site. Estimates of the BF at the LSO injection site are 14.8 kHz (A), and 10.25 kHz (B). *Top*, the response maps of ICC type I units with best frequencies below; *middle*, units with similar BFs; and *bottom*, units with BFs above the estimated BF at the injection site in the LSO. Responses are shown before (light gray) and during (dark gray) injection.

A second potential explanation could be the presence of an excitatory input derived from a non-LSO source. In this possibility, monaural and binaural response characteristics may differ between significantly affected and unaffected type I units due to the differing response characteristics of their inputs (i.e. LSO in the former and a non-LSO source in the latter). Figure 10A shows example BF tone rate-level curves for two type I units, one significantly affected (black) and one unaffected (gray). Driven discharge rates are shown as solid lines, spontaneous discharge rates as dashed lines. These units, which are representative of the populations of affected and unaffected units (summary statistics included in Table 2), respond with similar response characteristics (albeit slightly shifted in rate and level), including spontaneous rate (dashed lines), BF-tone threshold, maximum discharge rate, and plateau slope, and are overall comparable to values reported for type I units previously (Greene et al. 2010; Ramachandran et al. 1999). Example frequency response maps at 10 and 40 dB above threshold from two different type I units, one significantly affected (black) and one unaffected (gray), are shown normalized by BF in Fig. 10B. Both units show narrow V- or I-shaped excitatory tuning with flanking inhibition characteristic of type I unit responses (Ramachandran et al. 1999), and consistent with the comparable tuning (quantified as Q_10_ and Q_40_) recorded in the two populations (Table 2). The response map characteristics of affected and unaffected units, therefore, show no clear differences in these monaural response characteristics.

**Fig. 10.**
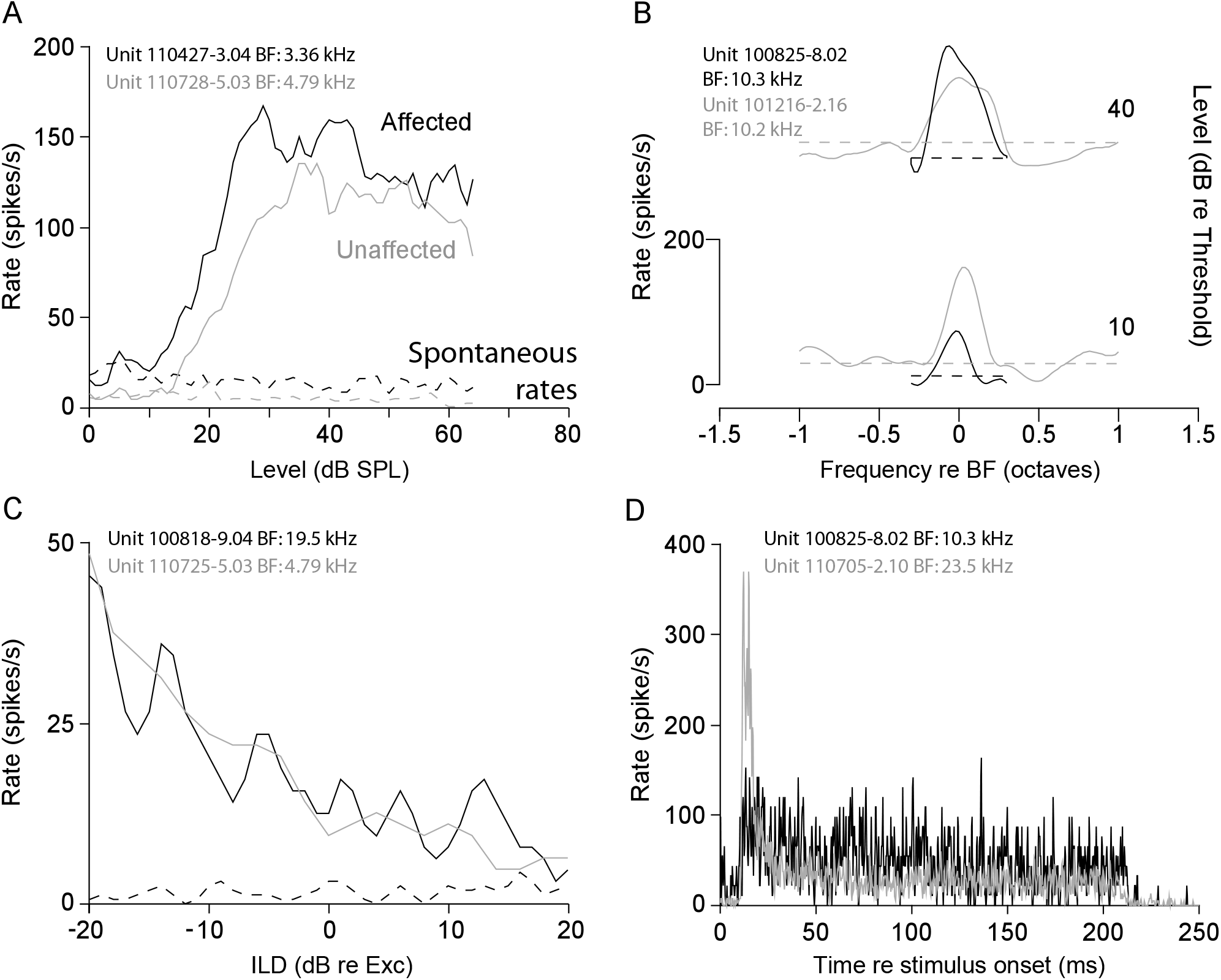
The response characteristics of affected (black) and unaffected (gray) type I units are largely comparable. The discharge rates of example affected and unaffected units, with relatively closely matched response characteristics, are shown in response to: A: BF tone rate-level curves; B: frequency tuning functions at 10 dB and 40 dB above threshold; C: interaural level difference functions recorded at approximately 10 dB re threshold; and D: BF tone post-stimulus time histograms recorded at approximately 20 dB re threshold.

**Table 2:**
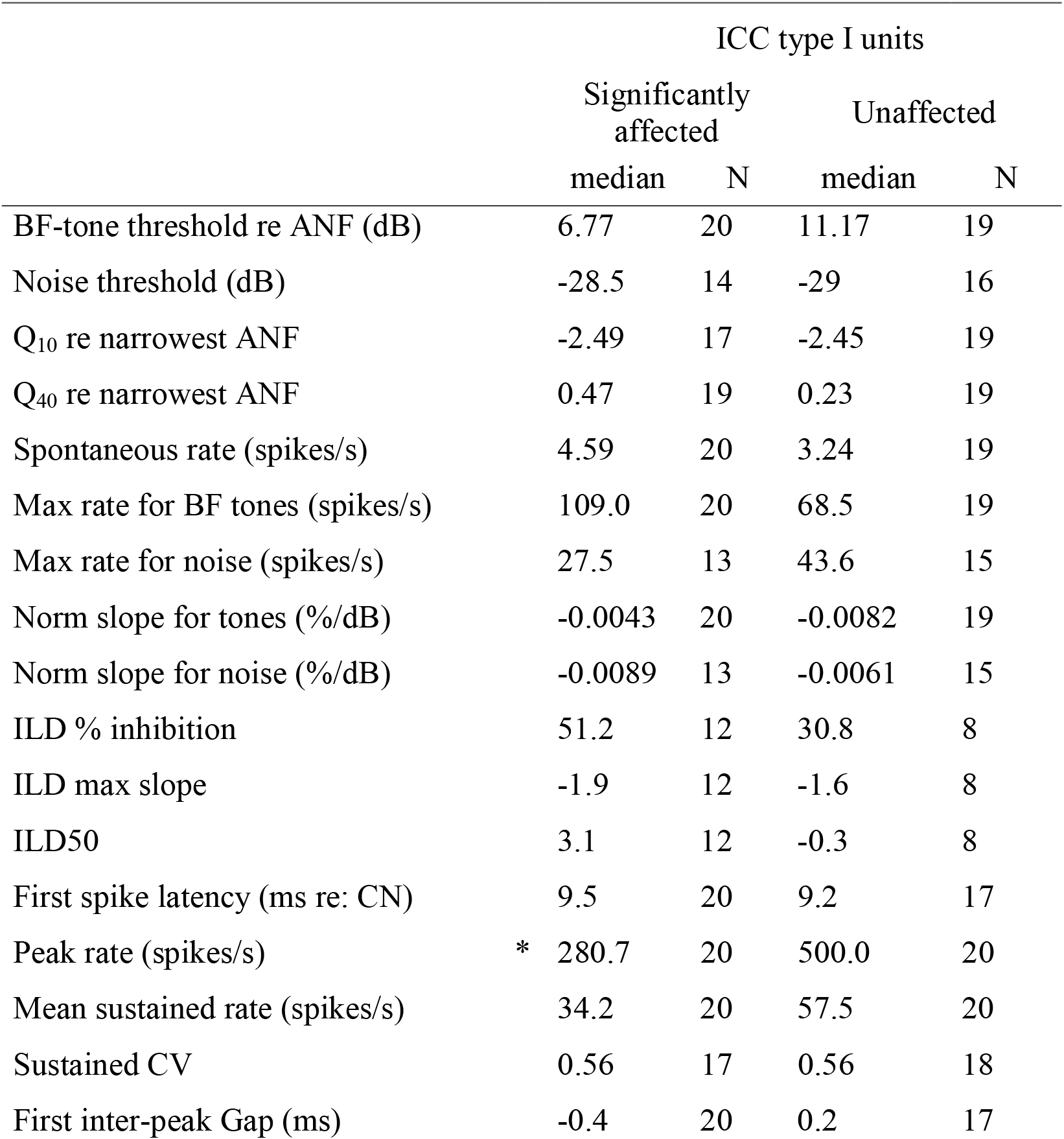
Baseline responses characteristics of affected and unaffected type I units. Values are medians across the populations of each group of type I units. Parameters that show a significant difference (Mann-whitney U test, p < 0.05) are marked with a *.

|  | ICC type I units |  |  |  |
| --- | --- | --- | --- | --- |
|  | Significantly affected |  | Unaffected |  |
|  | median | N | median | N |
| BF-tone threshold re ANF (dB) | 6.77 | 20 | 11.17 | 19 |
| Noise threshold (dB) | -28.5 | 14 | -29 | 16 |
| Q<sub>10</sub> re narrowest ANF | -2.49 | 17 | -2.45 | 19 |
| Q<sub>40</sub> re narrowest ANF | 0.47 | 19 | 0.23 | 19 |
| Spontaneous rate (spikes/s) | 4.59 | 20 | 3.24 | 19 |
| Max rate for BF tones (spikes/s) | 109.0 | 20 | 68.5 | 19 |
| Max rate for noise (spikes/s) | 27.5 | 13 | 43.6 | 15 |
| Norm slope for tones (%/dB) | -0.0043 | 20 | -0.0082 | 19 |
| Norm slope for noise (%/dB) | -0.0089 | 13 | -0.0061 | 15 |
| ILD % inhibition | 51.2 | 12 | 30.8 | 8 |
| ILD max slope | -1.9 | 12 | -1.6 | 8 |
| ILD50 | 3.1 | 12 | -0.3 | 8 |
| First spike latency (ms re: CN) | 9.5 | 20 | 9.2 | 17 |
| Peak rate (spikes/s) | * 280.7 | 20 | 500.0 | 20 |
| Mean sustained rate (spikes/s) | 34.2 | 20 | 57.5 | 20 |
| Sustained CV | 0.56 | 17 | 0.56 | 18 |
| First inter-peak Gap (ms) | -0.4 | 20 | 0.2 | 17 |

If the unaffected type I units receive their dominant excitatory input from a source other than LSO, we might expect substantial differences in two responses in particular. First, since LSO is generally considered to initiate an ILD processing pathway, we might expect to observe a difference in binaural sensitivity. Example responses from significantly affected (black) and unaffected (gray) type I units are shown in Fig. 10C, which show their driven (solid) and spontaneous (dashed) discharge rates as a function of ILD. As represented by these example responses, the half-maximal points, percentage inhibition, and normalized slopes (at the half-maximal point) were not different across the two populations (Table 2; note that these examples were chosen to highlight response similarity, and the differences is response variability are not observed at the population level).

Second, input from a different pathway may result in differing spike discharge characteristics in unaffected than affected type I units. The post-stimulus time (PST) histograms of two example type I units, showing discharge patterns representative of significantly affected (black) and unaffected (gray) type I units are shown in figure 10D. Since the responses to repeated tone stimulation were collected in order to assess the effects and time-course of LSO inactivation (e.g. Figs. 2–5), it was possible to generate post-stimulus time (PST) histograms by collecting these responses before response onset (R_ON_; Fig. 2). Several discharge pattern characteristics previously assessed in decerebrate cat LSO (Greene et al. 2012) were compared between affected and unaffected type I unit. Although the PSTs of unaffected units show onset peaks (i.e. the transient response at the start of the tone driven response) with significantly (Wann-Whitney U test, p < 0.05) higher instantaneous discharge rates on average than significantly affected type I units, other discharge characteristics are mostly comparable: minimum first spike latency, mean sustained (defined as the response during the last 80% of the stimulus) discharge rate, and the sustained coefficient of variation (CV; a measure of spike timing regularity) are not significantly different (summary statistics available in Table 2).

For the sake of brevity, we show only example units in Figure 10 and summary statistics in Tables 1 and 2. A more complete description of type I unit response characteristics is provided. Figure 11A compares unit BFs as a function of BF-tone thresholds for significantly affected (black) and unaffected (gray) units to the lowest threshold recorded in the auditory nerve, which are comparable to prior reports in decerebrate cat (Greene et al. 2010; Ramachandran et al. 1999), and were not significantly different from one another (Table 2). Figure 11B compares BF-tone thresholds recorded during KYNA injection to baseline recordings for those units with sufficient remaining activity that a threshold could be estimated. Thresholds were generally comparable across the two conditions, though significantly affected type I units tended to show increased thresholds during KYNA injection.

**Fig. 11.**
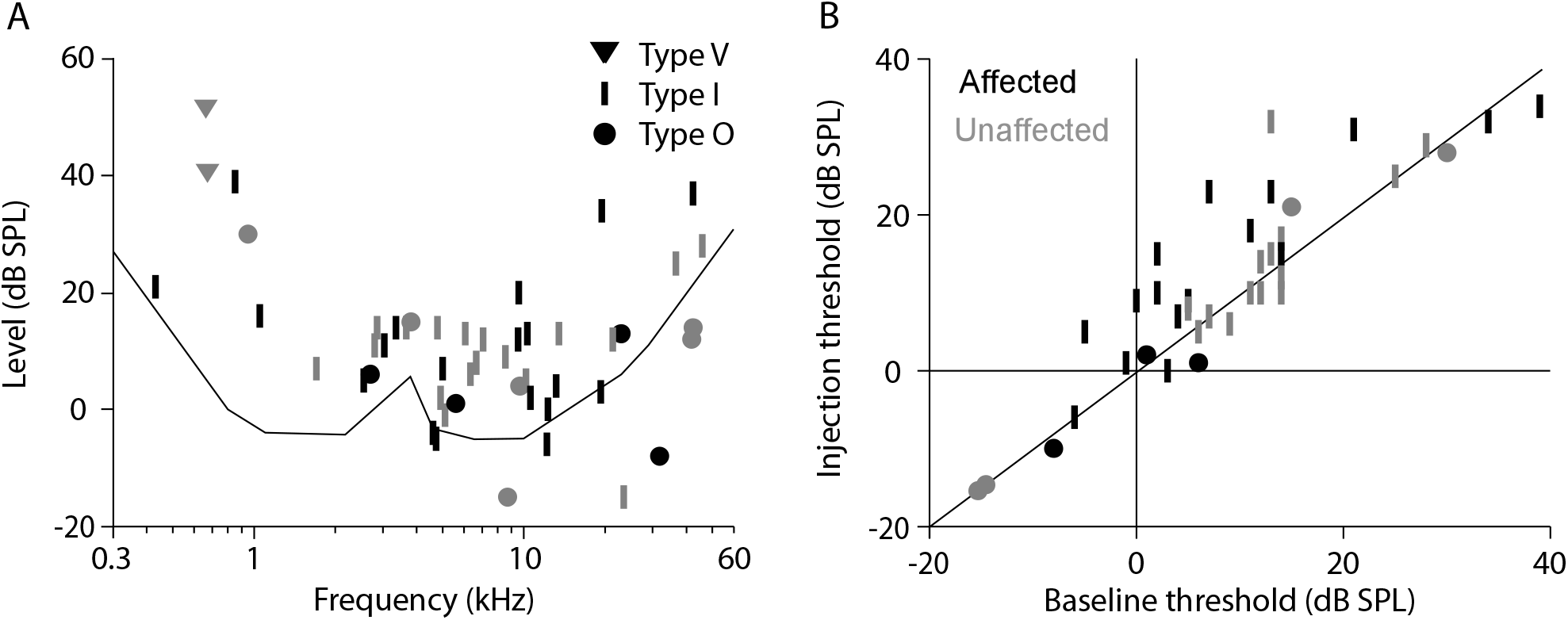
A: BF tone thresholds calculated from rate-level functions presented as a function of unit BF for all recorded ICC units. B: BF tone thresholds during the injection compared to baseline recordings for those units with sufficient residual activity to estimate threshold.

Figure 12A shows the example rate-level functions from Fig. 10A, along with straight line fits to the plateau region of the rate-level function (for a detailed description of the fit procedure, see: Greene et al. 2010). Fig. 12B-D show histograms of response characteristics across the populations of significantly affected and unaffected units, with the population median represented by a marker at the top of each plot: B) spontaneous discharge rate, C) maximum discharge rate (lowest intensity on the straight line fit), and D) the normalized slope of the straight line fit. No significant differences were noted in these rate-level function response characteristics (Table 2).

**Fig. 12.**
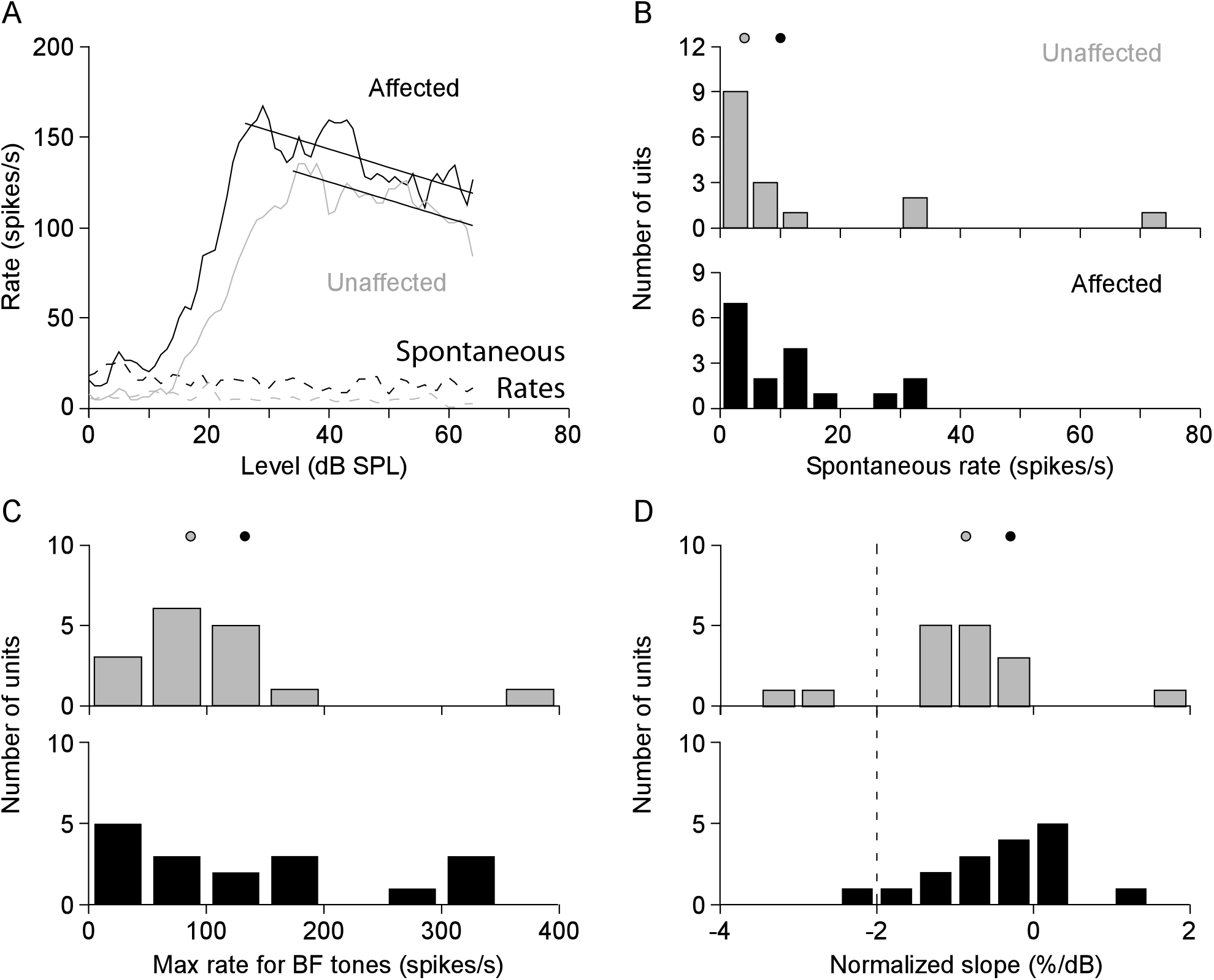
A: BF tone rate-level functions for two example units. Driven (solid) and spontaneous (dashed) discharge rates are shown, along with straight line fits to the plateau section (black lines). B: Histograms of spontaneous discharge rates in both unit populations. C: Histograms of the maximum discharge rate (defined as the point on the straight line fit at the lowest sound level) observed in the rate-level functions. D: Histograms of the slope of each straight line fit normalized to the maximum discharge rate observed for both populations of ICC type I units. Markers above each set of histograms indicate the medians of their respective populations.

Tuning curve width at 10 and 40 dB above threshold is compared between significantly and unaffected units in Figure 13, which shows the example units from Fig. 10B in Fig. 13A. The measured Q_10_ and Q_40_ values as a function of unit BF are superimposed onto lines delineating the range of responses observed in auditory nerve in Figs. 13B and C, and the difference in Q_40_ value between each unit and the highest value recorded in the auditory nerve (for a detailed description of Q value calculation and analysis, see: Greene et al. 2010). No significant differences were noted in tuning with across significantly affected and unaffected units (Table 2).

**Fig. 13.**
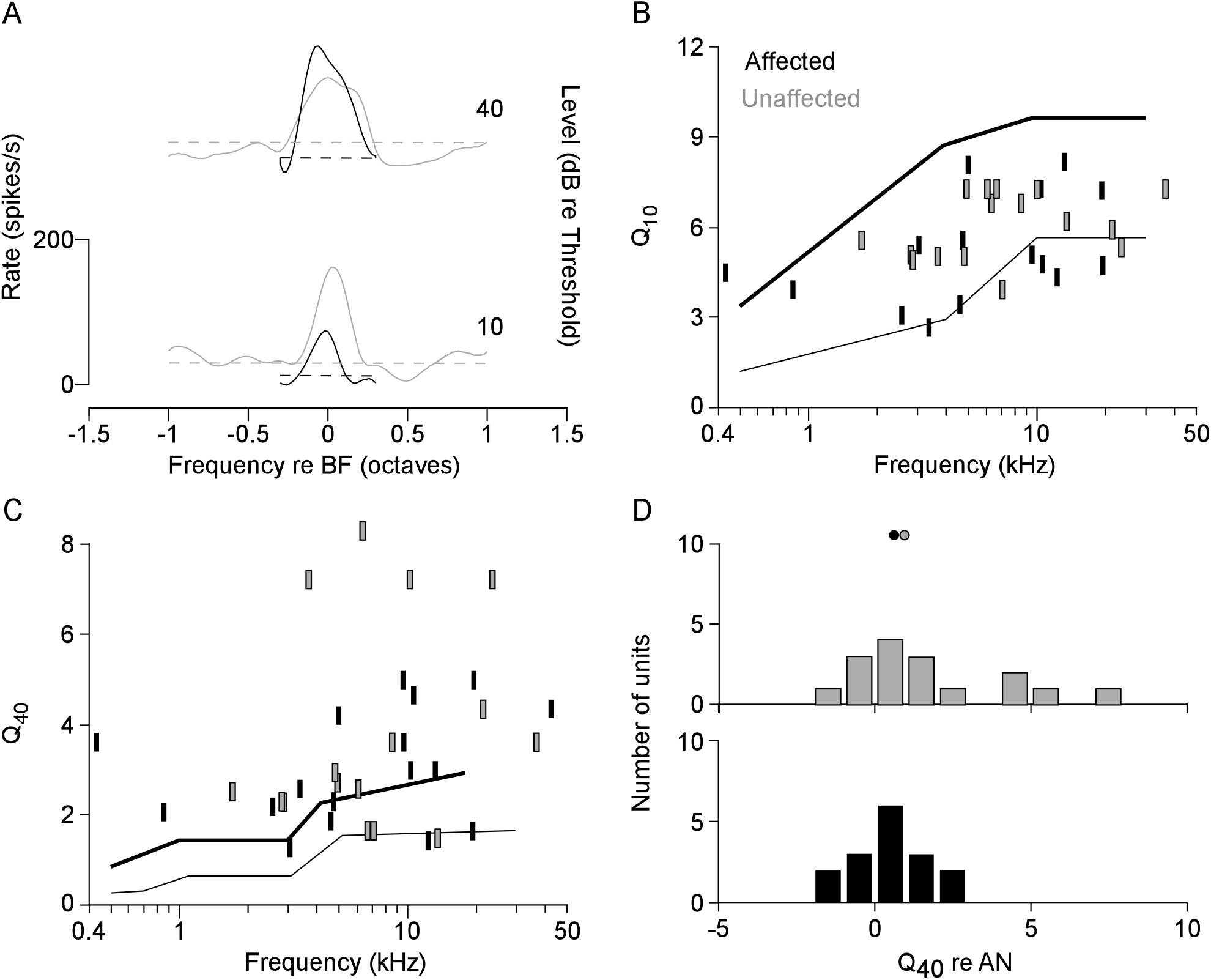
A: Frequency response maps of two example units measured at 10 and 40 dB above threshold. Driven discharge rates are indicated by line height, spontaneous discharge rates are shown with dashed lines. B: Q_10_ of each type I unit plotted as a function of unit BF, and C: Q_40_ of each type I unit plotted as a function of unit BF. B and C show unit tuning superimposed on the range of tuning observed in the auditory nerve. Black (gray) markers indicate significantly affected (unaffected) type I units. D: Histograms of the difference between type I unit tuning and the best tuning observed in the auditory nerve (upper line in C). Markers at the top indicate distribution medians.

ILD response characteristics are compared in Figure 14. The example responses from Fig. 10C are replicated in Fig. 14A. Sigmoidal fits to the rate-ILD curves are shown as smooth solid lines. The range of percentage inhibitions (calculated from the decrease of the sigmoidal fit between +20 and −20 dB ILD) observed in the significantly affected and unaffected populations of units are shown in Fig. 14B. Likewise, the range of ILD 50 point (defined as the ILD halfway between the upper and lower asymptotes in the sigmoid fits), marked in Fig. 14A with a small vertical tick mark, are shown in Fig. 14C. The range of maximum slopes observed in the sigmoid fits (the slope of the fit at the ILD 50 point) are shown in Fig. 14D. No significant differences were noted between the populations of significantly affected and unaffected units (Table 2).

**Fig. 14.**
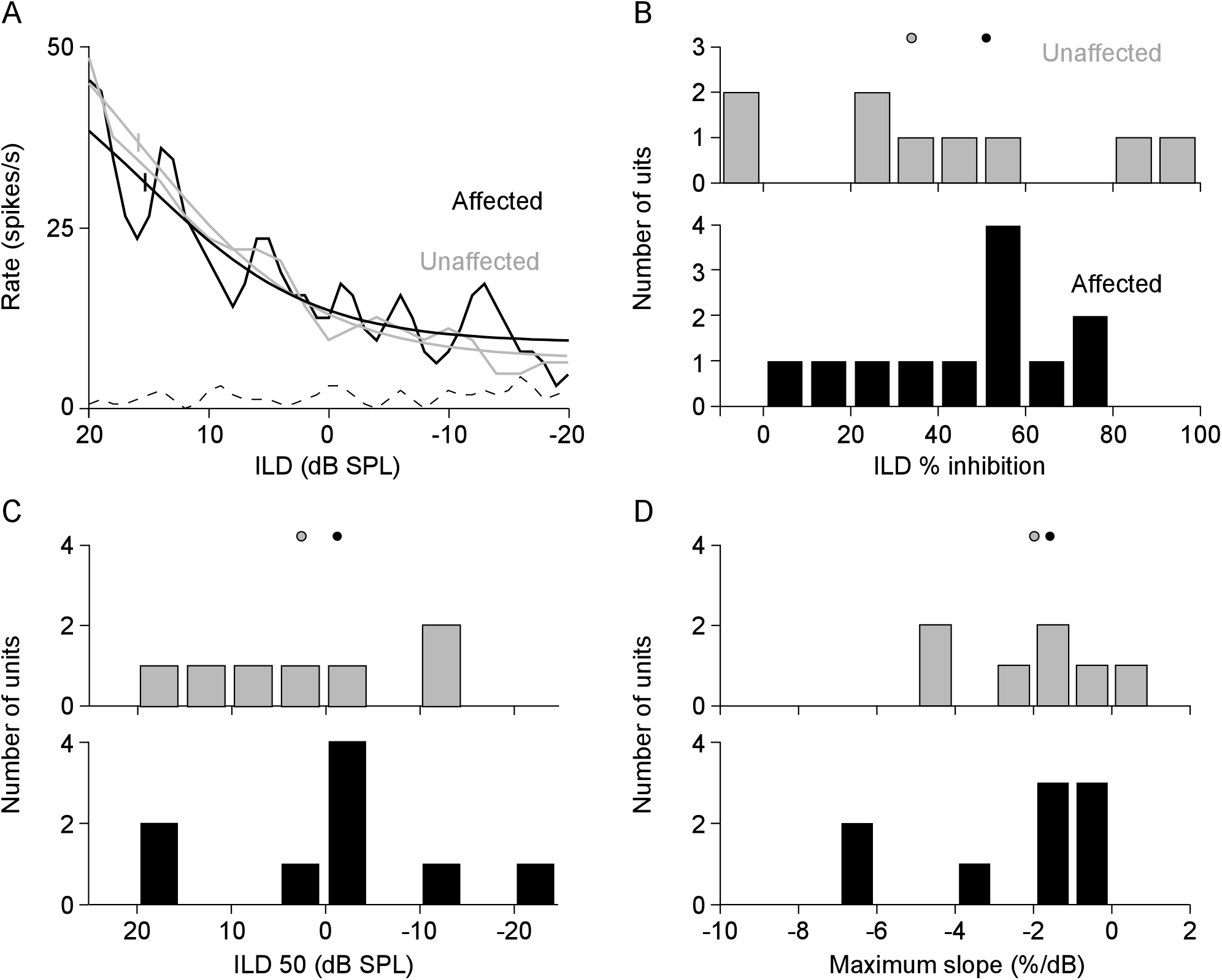
A: ILD functions observed in two example type I units. Driven discharge rates are indicated with solid lines, spontaneous rates with dashed lines (the spontaneous rate was zero during injection). Sigmoid fit lines are superimposed on these driven rate curves. Vertical tick marks indicate the ILD 50 point (the ILD halfway between the upper and lower asymptotes). B: Histograms of the degree of inhibition observed in each unit population (shown as a % change observed in the fit lines between +20 and −20 dB ILD). C: Histrograms of the ILD 50 points in the two populations of type I units. D: Histograms of the maximum slope (i.e. the slope at the ILD 50 point) observed in each sigmoidal fit line. Markers above each set of histograms indicate the median of each distribution.

Figures 15A and 1B shows the example PST histograms from Fig. 10D. The minimum first spike latency as a function of BF for all type I units is shown in Fig. 15C along with a line indicating the shortest latencies observed in cochlear nucleus (Young et al. 1988). The number of spikes recorded in the second peak as a function of the number of spikes in the first peak of the response (both normalized to the number of trials, thus a 1 indicates that a spike occurred in that peak on each trial) is shown in Fig. 15D. Most response characteristics tested were not significantly different between the significantly affected and unaffected units; however, unaffected units showed a significantly higher initial peak rate than significantly affected units, thus the spikes per peak values were significantly higher (for a detailed description of the analysis techniques, see: Greene and Davis 2012).

**Fig. 15.**
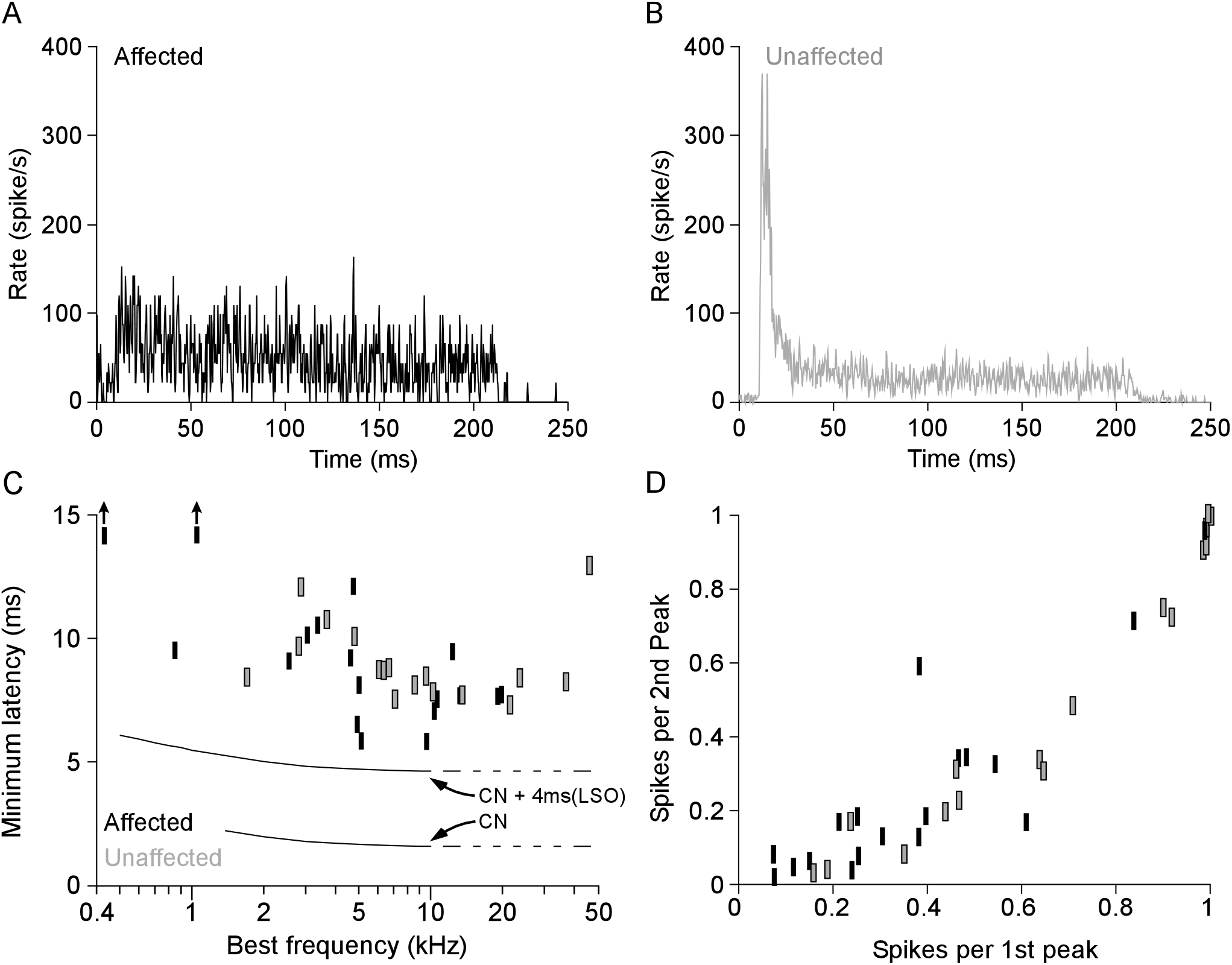
Example PST histograms recorded in A: a significantly affected, and B: an unaffected ICC type I unit. C: Minimim first spike latency of each type I unit as a function of unit BF, along with a line indicating the minimum first spike latency observed in the cochlear nucleus (as well as this line shifted + 4 ms) (Young et al. 1988). D: The number of spikes (normalized to the number of trial repetitions) observed in the second peak, as a function of the number of spikes observed in the first peak of the response. Black (gray) markers indicate significantly affected (unaffected) type I unit responses (for a detailed description of these analyses, see: Greene and Davis 2012).

## Discussion

The data presented above provide direct evidence that ICC type I units selectively receive direct, dominant excitatory input from the contralateral LSO. The responses of affected type I units can be dramatically reduced or silenced following injection of the non-specific excitatory amino antagonist kynurenic acid directly into LSO. These results are consistent with the excitatory projections of LSO forming a functionally segregated pathway through type I units in the ICC.

### Specificity of the LSO projection

Evidence for segregation of LSO projections in the ICC is principally provided by the scarcity of significantly and strongly affected type V or O units. In particular, the set of units in Fig. 4, where the I, but neither the V nor the O, type unit was strongly affected provides a test with single-units that were as similar as possible. That the LSO unit was recorded simultaneously provides direct evidence of the drug’s effect. However, few type V units were recorded, and those present showed somewhat differing effects, thus we are unable to draw any firm conclusions regarding this unit population. Nevertheless, it is unlikely that LSO contributes a substantial input to these cells since the excitatory projections of LSO are largely high frequency, while the BFs of type V units are largely low frequency. Type I and O units, on the other hand, both show high frequency tuning, thus could both show substantial LSO input. Inactivation of the DCN output tract (the dorsal acoustic stria) with lidocaine reveals that most (~80%) type O units receive their dominant excitatory input from DCN, thus a substantial contribution from LSO appears unlikely (Davis 2002). The current recordings were biased against type V and O units for methodological reasons (a focus on high-frequency units, and an attempt to record a strongly affected unit early in the experiment to prove that the drug was effective and the electrode was functional); however, for the reasons stated above, we do not expect the effects of LSO inactivation to strongly affect the majority of these units. Nevertheless, the type O unit responses remaining following DAS blackade are created de novo in the ICC, thus we cannot rule out altogether a LSO contribution.

Results from physiological comparisons (Greene et al. 2010) suggest that the projection from LSO to ICC type I units is a mostly high-frequency pathway, and since this projection is largely excitatory (Glendenning et al. 1992), the effect of silencing LSO activity should largely be a reduction in discharge rate. Consistent with this excitatatory input, most units affected by injection of kynurenic acid into LSO do indeed show a strong reduction in discharge rate. However, we do not observe the BF dependence expected from the frequency bias in the projection, instead altered responses were observed in units with BFs spanning the entire recorded range. Although the projection from LSO to contralateral ICC is largely excitatory, direct inhibitory projections have been observed (Glendenning et al. 1992), and several indirect inhibitory pathways exist between LSO and contralateral ICC, particularly through ipsi- and conta-lateral DNLL, as well as contralateral ICC (Hernandez et al. 2006; Kelly and Li 1997; Zhang et al. 1998). In contrast to the direct excitatory effects, units receiving inhibitory input from LSO (either directly or indirectly) should show response rates that generally increase during LSO inactivation. Similarly an inhibitory input from contralateral LSO should decrease responses at low ILDs, thus an ICC unit with an E) binaural response should show an increasing rate with ILD, producing an ILD response consistent with an EE response. A small minority of units (two type I and one type V) recorded showed slight but significant increases in discharge rate during KYNA injection, which suggests that these inhibitory inputs can influence ICC unit responses. Definitive tests of both the specificity of the low frequency projection and effects of direct and indirect inhibitory pathways from LSO to ICC will require more thorough tests of LSO projection patterns.

Over half of type I unit’s tone driven activity is significantly reduced by pharmacological inactivation of the contralateral LSO. For a number of reasons, however, our estimate of the proportion of LSO derived type I units is a lower bound on the total number. First, the location of LSO is deep within the brainstem, thus limiting the type of electrodes available (as electrode width becomes increasingly important), as well as increasing the risk of insertion errors, making incomplete LSO inactivation more likely. Use of the carbon fiber electrodes (and insulated hypodermic needle) minimized damage to overlying brainstem structures compared to a piggy-back glass micropipette/metal electrode, but prevented isolation of single units (except in a minority of cases: Fig. 2, 4), and required the use a two-step insertion procedure increasing the chance of insertion errors. Second, since the significance of the LSO inactivation effect in ICC unit responses was defined at a single suprathreshold level at BF, the magnitude of effect following LSO inactivation is underestimated for units showing evidence for convergence with non-monotonic (with intensity) projections such as those from DCN (e.g. Fig. 8), or other unexpected changes. Third, the method used to objectively identify R_ON_ and R_SS_ often provides a conservative estimate of these measures (e.g. Fig. 3B R_ON_ appears later than would be placed by eye), thus the magnitude of the KYNA effect is somewhat decreased in some units (since some decrease is included in the baseline rate estimate). Finally, the similarities between significantly affected and unaffected unit populations, as well as differing effects in units with similar locations in ICC or with similar response properties (e.g. Fig. 9) in individual experiments, could be evidence for incomplete LSO suppression.

### On the origins of unaffected type I units

A large proportion of type I units reported here show no indication of excitatory input from LSO. There are two ways by which type I units could show no effect in the current study. First, kynurenic acid injections may have been insufficiently large to silence LSO activity completely. The LSO is a relatively large 3-dimensional structure, with a highly ordered internal structure (Cant 1984). Drug diffusion, therefore, may not be perfectly spherical, and instead may preferentially spread either within an isofrequency contour, or along the tonotopic gradient. Indeed, an example of drug favoring diffusion along the length of LSO cells (i.e. within an isofrequency contour) is apparent following HRP injection in cat (Fig. 3 from Glendenning et al. 1985). Anecdotal evidence assessing the possibility for anisotropic spread of the drug is shown here by the two seemingly contradictory example electrode penetrations shown in Fig. 9. In the first (Fig. 9A), kynurenic acid injection appeared to only affect units with a BF similar to the site of injection within the LSO. In the second (Fig. 9B), kynurenic acid injections spared units with similar BFs, but strongly affected units with higher and lower BFs, suggesting injection incompletely inactivated LSO units within the isofrequency contour. A more definitive assessment of this possibility will require methodology with greater spatial selectivity.

The second mechanism by which type I units could avoid the influence of LSO input is by way of non-LSO excitatory inputs. Excitatory inputs may derive from other, non-LSO, brainstem regions; in particular, ventral cochlear nucleus multipolar cells provide a large source of compatible excitatory drive to contralateral ICC (e.g. Cant and Benson 2003). In this case, inhibitory inputs must produce binaural sensitivity within the ICC, which could derive from a number of brainstem nuclei including areas dominated by glycinergic neurons such as VNLL and ipsilateral LSO, as well as GABAergic neurons such as DNLL or contralateral ICC (Batra and Fitzpatrick 2002; Glendenning et al. 1992; Kelly and Li 1997; Malmierca et al. 2003). In a direct test of the magnitude of inhibitory effect via pharmacological blockade in the bat, one study suggested that ILD sensitivity is modulated in the ICC in as many as half of EI sensitive ICC units (Klug et al. 1995). Glycinergic blockade affected ipsilateral suppression in one-third of units, but eliminated ILD sensitivity in only 8% (5/61) of units; GABAergic blockade was comparable, with ipsilateral inhibition suppressed in 40% (15/38) and eliminated in 13% (5/38) of units. Moderately affected units could still derive excitatory input from LSO, suggesting that no more than ~20% of EI units in the bat are derived from inputs other than LSO. Furthermore, the large number of EI units moderately affected by inhibitory blockade is perhaps not surprising, as several studies have previously suggested that inhibitory inputs could produce hierarchical transformations, thus increasing response complexity in ICC cells compared to LSO (Greene et al. 2010; Nelson and Carney 2004; Park et al. 1996; Park and Pollak 1993; Tollin and Yin 2002a; b; Tsai et al. 2010).

Type I unit responses that receive non-LSO excitatory input would likely have different response characteristics, consistent with their different inputs. In particular, if responses are derived from ventral cochlear nucleus, we might expect to observe shorter first spike latency consistent with a more direct projection pathway, different frequency tuning and level responses, and weaker ILD sensitivity. Direct comparisons of response characteristics in Fig. 10 and table 2, however, suggest that units affected and unaffected by LSO inactivation show largely comparable response characteristics. Indeed, only one property varies in any substantial manner: discharge patterns indicate more prominent onset peaks in unaffected units. This could indicate a greater influence of a delayed inhibitory input in unaffected than in affected units, but the differences are relatively small and after correcting for [the large number of] multiple comparisons, not significant. Comparisons between response characteristics, therefore, suggest that responses either derive from cells with similar response characteristics, or that inhibitory inputs act to produce nearly identical responses in the ICC. One important caveat to this result, however, is that comparisons are only presented for relatively simple stimuli. Greater differences between these units may appear in the responses to stimuli for which hierarchical processing is expected, such as in the processing of ILD as a function of level (Greene et al. 2008; Park 1998; Park et al. 2004; Tsai et al. 2010), or in the formation of EI/f ILD responses in the ICC (Davis et al. 1999; Park and Pollak 1993).

### Origins of ILD sensitivity in the ICC and the role of the LSO

The similarities in monaural and binaural response properties between LSO and ICC type I units form the basis for the hypothesis that LSO projections are the dominant excitatory input to all type I units in the ICC, and that this projection initiates a pathway specialized to process ILD cues (Davis et al. 1999; Greene et al. 2010; Ramachandran et al. 1999). Supporting this hypothesis, behavioral studies following (irreversible) kainic acid lesions have demonstrated the importance of the superior olive to sound localization (Kavanagh and Kelly 1992; van Adel and Kelly 1998). However, single cell recordings during similar experiments show that ILD sensitive units remain in the ICC following superior olive lesion (Kelly and Sally 1993; Sally and Kelly 1992), and intracellular recordings in the ICC reveal de novo creation of ILD sensitivity in the ICC that is not inherited from LSO (Li et al. 2010; Xie et al. 2008), suggesting that ILD sensitivity is not solely dependent upon olivary input.

Methodological and species differences suggest that the results of these prior studies do not address the hypothesis that LSO initiates, and that ICC type I units are the midbrain component of, a functionally segregated pathway specialized to process ILD: in particular, the responses of high-frequency units in decerebrate cat differ in several ways from anesthetized rat. First, in decerebrate cat, both type I and type O units show EI binaural response characteristics, but type O units, like their DCN principal cell inputs, are distinguishable based upon the large amount of suppression observed at high sound levels (Davis et al. 1999). In contrast, responses of DCN principal cells in rodents produce type III responses more frequently (which show no such high level inhibition), and type IV responses less frequently than in cat (gerbil: Davis et al. 1996), (rat: Yajima and Hayashi 1989), (mouse: Ma and Brenowitz 2012). The similarity between rodent DCN and LSO unit monaural response maps suggests that ICC units receiving DCN input in rodents will show response maps more similar to type I than type O units, potentially confounding those results. Second, anesthesia is known to alter the balance of excitation and inhibition in the auditory brainstem. In particular, the dominant response type of the DCN principal cells in the cat shifts from type IV to I/III under the influence of barbiturate anesthesia (Evans and Nelson 1973), further decreasing the distinction between LSO and DCN derived inputs in ICC unit responses. Projections of DCN to ICC in anesthetized rat may thus be difficult to distinguish from the projections of LSO based on response map characteristics, which may explain at least some of the remaining ILD sensitivity observed in ICC units following kainic acid lesion of the SOC (Kelly and Sally 1993; Sally and Kelly 1992).

Nevertheless, these previous studies can help provide context to the current results. The kainic acid lesion studies suggest that LSO plays a critical role in sound source localization localization (Kavanagh and Kelly 1992; van Adel and Kelly 1998), despite the existence of remaining ILD sensitivity in ICC unit cells (Kelly and Sally 1993; Sally and Kelly 1992). The current results suggest that the activity of most, though probably not all (Klug et al. 1995), ILD sensitive cells in ICC is derived in contralateral LSO. Thus, since the binaural responses of those ILD sensitive cells remaining after SOC ablation must be a result of input from other binaural nuclei (e.g. DCN) or de-novo bilateral inputs within the ICC, the binaurality of these remaining units is likely a side-effect of process some other stimulus characteristic. Conversely, since the LSO appears to be critical for sound source localization using ILDs, and the dominant excitatory input to ICC type I units appears to derive from contralateral LSO, ILD process appears to occur within a functionally segregated pathway at least through the auditory midbrain. Additional investigation is required to determine whether this pathway extends to higher levels within the auditory system (i.e. thalamus or cortex), whether additional ILD processing occurs downstream, and whether this pathway is involved in the processing of additional auditory features or ILD alone.

## Acknowledgements

We are grateful for the helpful input of Drs. L. Carney, W. O’Neill, G. Paige, and D. Tollin, as well as technical assistance from O. Lomakin and T. Bubel.

## References

Aitkin LM, Webster WR, Veale JL, and Crosby DC. Inferior colliculus. I. Comparison of response properties of neurons in central, pericentral, and external nuclei of adult cat. J Neurophysiol 38: 1196–1207, 1975.

Batra R, and Fitzpatrick DC. Monaural and binaural processing in the ventral nucleus of the lateral lemniscus: a major source of inhibition to the inferior colliculus. Hear Res 168: 90–97, 2002.

Brownell WE, Manis PB, and Ritz LA. Ipsilateral inhibitory responses in the cat lateral superior olive. Brain Res 177: 189–193, 1979.

Burger RM, and Pollak GD. Reversible inactivation of the dorsal nucleus of the lateral lemniscus reveals its role in the processing of multiple sound sources in the inferior colliculus of bats. J Neurosci 21: 4830–4843, 2001.

Caird D, and Klinke R. Processing of binaural stimuli by cat superior olivary complex neurons. Exp Brain Res 52: 385–399, 1983.

Cant NB. The fine structure of the lateral superior olivary nucleus of the cat. In: Journal of comparative neurology. Philadelphia, Pa.: Wistar Institute of Anatomy and Biology, 1984, p. 63.

Cant NB, and Benson CG. Parallel auditory pathways: projection patterns of the different neuronal populations in the dorsal and ventral cochlear nuclei. Brain Res Bull 60: 457–474, 2003.

Davis KA. The basic receptive field properties of the neurons in the inferior colliculus of decerebrated cats are rarely created by local inhibitory mechanisms. In: Soc Neurosci Abstr 1999, p. 667.

Davis KA. Contralateral effects and binaural interactions in dorsal cochlear nucleus. J Assoc Res Otolaryngol 6: 280–296, 2005.

Davis KA. Evidence of a functionally segregated pathway from dorsal cochlear nucleus to inferior colliculus. J Neurophysiol 87: 1824–1835, 2002.

Davis KA, Ding J, Benson TE, and Voigt HF. Response properties of units in the dorsal cochlear nucleus of unanesthetized decerebrate gerbil. J Neurophysiol 75: 1411–1431, 1996.

Davis KA, Ramachandran R, and May BJ. Single-unit responses in the inferior colliculus of decerebrate cats. II. Sensitivity to interaural level differences. J Neurophysiol 82: 164–175, 1999.

Evans EF, and Nelson PG. The responses of single neurones in the cochlear nucleus of the cat as a function of their location and the anaesthetic state. Exp Brain Res 17: 402–427, 1973.

Faingold CL, Anderson CA, and Randall ME. Stimulation or blockade of the dorsal nucleus of the lateral lemniscus alters binaural and tonic inhibition in contralateral inferior colliculus neurons. Hear Res 69: 98–106, 1993.

Finlayson PG, and Caspary DM. Low-frequency neurons in the lateral superior olive exhibit phase-sensitive binaural inhibition. J Neurophysiol 65: 598–605, 1991.

Gibson DJ. Interaural crosstalk in the cat. Hear Res 7: 325–333, 1982.

Glendenning KK, Baker BN, Hutson KA, and Masterton RB. Acoustic chiasm V: inhibition and excitation in the ipsilateral and contralateral projections of LSO. J Comp Neurol 319: 100–122, 1992.

Glendenning KK, Hutson KA, Nudo RJ, and Masterton RB. Acoustic chiasm II: Anatomical basis of binaurality in lateral superior olive of cat. J Comp Neurol 232: 261–285, 1985.

Goldberg JM, and Brown PB. Response of binaural neurons of dog superior olivary complex to dichotic tonal stimuli: some physiological mechanisms of sound localization. J Neurophysiol 32: 613–636, 1969.

Greene NT, and Davis KA. Discharge patterns in the lateral superior olive of decerebrate cats. J Neurophysiol 108: 1942–1953, 2012.

Greene NT, Lomakin O, and Davis KA. Monaural spectral processing differs between the lateral superior olive and the inferior colliculus: physiological evidence for an acoustic chiasm. Hear Res 269: 134–145, 2010.

Greene NT, Lomakin OA, and Davis KA. Response Properties of Single Units in the Lateral Superior Olive of Decerebrate Cats. In: Assoc Res Otolaryngol Abs/2008, p. 855.

Guinan JJ, Jr., Guinan SS, and Norris BE. Single auditory units in the superior olivary complex. I. Responses to sounds and classifications based on physiological properties. Int J Neurosci 4: 101–120, 1972a.

Guinan JJ, Jr., Norris BE, and Guinan SS. Single auditory units in the superior olivary complex. II. Locations of unit categories and tonotopic organization. Int J Neurosci 4: 147–166, 1972b.

Hernandez O, Rees A, and Malmierca MS. A GABAergic component in the commissure of the inferior colliculus in rat. Neuroreport 17: 1611–1614, 2006.

Kavanagh GL, and Kelly JB. Midline and lateral field sound localization in the ferret (Mustela putorius): contribution of the superior olivary complex. J Neurophysiol 67: 1643–1658, 1992.

Kelly JB, and Li L. Two sources of inhibition affecting binaural evoked responses in the rat’s inferior colliculus: the dorsal nucleus of the lateral lemniscus and the superior olivary complex. Hear Res 104: 112–126, 1997.

Kelly JB, and Sally SL. Effects of superior olivary complex lesions on binaural responses in rat auditory cortex. Brain Res 605: 237–250, 1993.

Klug A, Park TJ, and Pollak GD. Glycine and GABA influence binaural processing in the inferior colliculus of the mustache bat. J Neurophysiol 74: 1701–1713, 1995.

LeBeau FE, Malmierca MS, and Rees A. Iontophoresis in vivo demonstrates a key role for GABA(A) and glycinergic inhibition in shaping frequency response areas in the inferior colliculus of guinea pig. J Neurosci 21: 7303–7312, 2001.

Li L, and Kelly JB. Binaural responses in rat inferior colliculus following kainic acid lesions of the superior olive: interaural intensity difference functions. Hear Res 61: 73–85, 1992a.

Li L, and Kelly JB. Inhibitory influence of the dorsal nucleus of the lateral lemniscus on binaural responses in the rat’s inferior colliculus. J Neurosci 12: 4530–4539, 1992b.

Li N, Gittelman JX, and Pollak GD. Intracellular recordings reveal novel features of neurons that code interaural intensity disparities in the inferior colliculus. J Neurosci 30: 14573–14584, 2010.

Ma WL, and Brenowitz SD. Single-neuron recordings from unanesthetized mouse dorsal cochlear nucleus. J Neurophysiol 107: 824–835, 2012.

Malmierca MS, Hernandez O, Falconi A, Lopez-Poveda EA, Merchan M, and Rees A. The commissure of the inferior colliculus shapes frequency response areas in rat: an in vivo study using reversible blockade with microinjection of kynurenic acid. Exp Brain Res 153: 522–529, 2003.

Merzenich MM, and Reid MD. Representation of the cochlea within the inferior colliculus of the cat. Brain Res 77: 397–415, 1974.

Nelson PC, and Carney LH. A phenomenological model of peripheral and central neural responses to amplitude-modulated tones. J Acoust Soc Am 116: 2173–2186, 2004.

Oliver DL, and Huerta MF. Inferior and superior colliculi. In: The mammalian auditory pathway, edited by Webster DB, Popper AN, and Fay RR. New York: Springer, 1992, p. 168–221.

Park TJ. IID sensitivity differs between two principal centers in the interaural intensity difference pathway: the LSO and the IC. J Neurophysiol 79: 2416–2431, 1998.

Park TJ, Grothe B, Pollak GD, Schuller G, and Koch U. Neural delays shape selectivity to interaural intensity differences in the lateral superior olive. J Neurosci 16: 6554–6566, 1996.

Park TJ, Klug A, Holinstat M, and Grothe B. Interaural level difference processing in the lateral superior olive and the inferior colliculus. J Neurophysiol 92: 289–301, 2004.

Park TJ, and Pollak GD. GABA shapes a topographic organization of response latency in the mustache bat’s inferior colliculus. J Neurosci 13: 5172–5187, 1993.

Ramachandran R, Davis KA, and May BJ. Single-unit responses in the inferior colliculus of decerebrate cats. I. Classification based on frequency response maps. J Neurophysiol 82: 152–163, 1999.

Ramachandran R, and May BJ. Functional segregation of ITD sensitivity in the inferior colliculus of decerebrate cats. J Neurophysiol 88: 2251–2261, 2002.

Rowland BA, Quessy S, Stanford TR, and Stein BE. Multisensory integration shortens physiological response latencies. J Neurosci 27: 5879–5884, 2007.

Sally SL, and Kelly JB. Effects of superior olivary complex lesions on binaural responses in rat inferior colliculus. Brain Res 572: 5–18, 1992.

Spirou GA, and Young ED. Organization of dorsal cochlear nucleus type IV unit response maps and their relationship to activation by bandlimited noise. J Neurophysiol 66: 1750–1768, 1991.

Suga N, O’Neill WE, and Manabe T. Cortical neurons sensitive to combinations of information-bearing elements of biosonar signals in the mustache bat. Science 200: 778–781, 1978.

Tollin DJ, and Koka K. Postnatal development of sound pressure transformations by the head and pinnae of the cat: Binaural characteristics. J Acoust Soc Am 126: 3125–3136, 2009a.

Tollin DJ, and Koka K. Postnatal development of sound pressure transformations by the head and pinnae of the cat: Monaural characteristics. J Acoust Soc Am 125: 980–994, 2009b.

Tollin DJ, Koka K, and Tsai JJ. Interaural level difference discrimination thresholds for single neurons in the lateral superior olive. J Neurosci 28: 4848–4860, 2008.

Tollin DJ, and Yin TC. The coding of spatial location by single units in the lateral superior olive of the cat. I. Spatial receptive fields in azimuth. J Neurosci 22: 1454–1467, 2002a.

Tollin DJ, and Yin TC. The coding of spatial location by single units in the lateral superior olive of the cat. II. The determinants of spatial receptive fields in azimuth. J Neurosci 22: 1468–1479, 2002b.

Tollin DJ, and Yin TC. Interaural phase and level difference sensitivity in low-frequency neurons in the lateral superior olive. J Neurosci 25: 10648–10657, 2005.

Tsai JJ, Koka K, and Tollin DJ. Varying overall sound intensity to the two ears impacts interaural level difference discrimination thresholds by single neurons in the lateral superior olive. J Neurophysiol 103: 875–886, 2010.

Tsuchitani C. Functional organization of lateral cell groups of cat superior olivary complex. J Neurophysiol 40: 296–318, 1977.

Tsuchitani C, and Boudreau JC. Single unit analysis of cat superior olive S segment with tonal stimuli. J Neurophysiol 29: 684–697, 1966.

van Adel BA, and Kelly JB. Kainic acid lesions of the superior olivary complex: effects on sound localization by the albino rat. Behav Neurosci 112: 432–446, 1998.

Vater M, Habbicht H, Kossl M, and Grothe B. The functional role of GABA and glycine in monaural and binaural processing in the inferior colliculus of horseshoe bats. J Comp Physiol A 171: 541–553, 1992.

Xie R, Gittelman JX, Li N, and Pollak GD. Whole cell recordings of intrinsic properties and sound-evoked responses from the inferior colliculus. Neuroscience 154: 245–256, 2008.

Xie R, Gittelman JX, and Pollak GD. Rethinking tuning: in vivo whole-cell recordings of the inferior colliculus in awake bats. J Neurosci 27: 9469–9481, 2007.

Yajima Y, and Hayashi Y. Response properties and tonotopical organization in the dorsal cochlear nucleus in rats. Exp Brain Res 75: 381–389, 1989.

Yang L, Pollak GD, and Resler C. GABAergic circuits sharpen tuning curves and modify response properties in the mustache bat inferior colliculus. J Neurophysiol 68: 1760–1774, 1992.

Yin TC, and Chan JC. Interaural time sensitivity in medial superior olive of cat. J Neurophysiol 64: 465–488, 1990.

Young ED, and Brownell WE. Responses to tones and noise of single cells in dorsal cochlear nucleus of unanesthetized cats. J Neurophysiol 39: 282–300, 1976.

Young ED, Robert JM, and Shofner WP. Regularity and latency of units in ventral cochlear nucleus: implications for unit classification and generation of response properties. J Neurophysiol 60: 1–29, 1988.

Zhang DX, Li L, Kelly JB, and Wu SH. GABAergic projections from the lateral lemniscus to the inferior colliculus of the rat. Hear Res 117: 1–12, 1998.

